# TCR signal strength determines T_reg_ instability and discrimination of self versus non-self antigens

**DOI:** 10.64898/2026.02.12.705537

**Authors:** Liping Li, Jie Guo, Ruiqiao He, Xin Chai, Chang Guo, Wei Xu, Xue Cao, Qiuzhu Jin, Fuping Zhang, Baidong Hou, Fangqing Zhao, Xuyu Zhou

## Abstract

Regulatory T cells (Tregs) expressing autoreactive T-cell receptors (TCRs) maintain immune self-tolerance, yet their fate upon antigen encounter remains unclear. Although selected in the thymus for high-affinity self-recognition, the cross-reactive nature of the TCR repertoire allows some Tregs to also recognize foreign antigens with high affinity. Here, we identify cross-reactive destabilization, a process in which high-affinity recognition of foreign antigens erodes Treg stability and redirects lineage fate. Using a thymic-derived Treg (tTreg) TCR capable of recognizing both self and foreign antigens distinct from the self-ligand that mediated its thymic selection, we show that stimulation with high-affinity non-self agonists induces Foxp3 loss and conversion into effector-like ex-Tregs, whereas self-antigen stimulation preserves lineage stability. Treg instability also increases with age and correlates with activation, suggesting that cumulative foreign-antigen exposure drives this process. These findings reveal a signal-strength–dependent model in which strong TCR engagement induces cross-reactive destabilization to balance immune responsiveness and self-tolerance.

**One Sentence Summary:** Foreign antigen-driven destabilization of T_reg_ cells balances immunity and tolerance

## INTRODUCTION

Regulatory T cells (T_reg_ cells) are central guardians of immune self-tolerance—their absence results in fatal autoimmune pathology(*1*). Foxp3, the master transcriptional regulator of T_reg_ identity, functions not merely as a lineage-specifying factor but as an essential molecular safeguard; its loss in humans (IPEX syndrome) and mice (scurfy) precipitates lethal inflammatory cascades, unequivocally demonstrating T_reg_ cells’ non-redundant role in immune homeostasis(*1–3*). Crucially, loss of Foxp3 expression in mature peripheral T_reg_ cells results in their complete reprogramming into effector-like cells, demonstrating that sustained Foxp3 expression is essential for maintaining T_reg_ stability and suppressive function(*4, 5*).

Like conventional CD4^+^ T cells (T_conv_ cells), T_reg_ cells express diverse T cell receptors (TCRs) capable of recognizing peptide-MHC class II complexes. TCR engagement by self-antigens is critical for T_reg_ cells to acquire suppressive competence and differentiate into specialized subsets, including T-bet^+^ (T_H_1-like) T_reg_ cells and Bcl6^+^ follicular regulatory T (T_FR_) cells (*6–9*). Paradoxically, these same TCR signals that confer functional specialization can also destabilize T_reg_ identity, driving Foxp3 downregulation and conversion into proinflammatory ex-T_reg_ cell (*10–12*). The molecular mechanisms distinguishing stabilizing from destabilizing TCR signals remain a fundamental unresolved question in immune regulation.

TCRs exhibit intrinsic cross-reactivity, enabling individual receptors to engage multiple peptide-MHC class II complexes (*13*). This property permits comprehensive immune surveillance despite the limited TCR repertoire (*14*). For T_reg_ cells—which frequently express autoreactive TCRs—this cross-reactivity creates a unique regulatory dilemma: while self-antigen-driven TCR signaling is essential for T_reg_ development and function, these same TCRs may exhibit varying responsiveness to foreign antigens.

T_FR_ cells present a fascinating immunological paradox: although resident in germinal centers (GCs), they display minimal responsiveness to the foreign antigens that initiate GC reactions. Clonal analyses demonstrate that T_FR_ cells are predominantly self-reactive, closely mirroring thymus-derived T_reg_ (tT_reg_) cells while showing negligible TCR repertoire overlap with T follicular helper (T_FH_) cells(*15*). Both TCR transgenic (Tg) mouse models and MHC class II tetramer studies consistently localize foreign antigen-specific T cells to the T_FH_ cell compartment, with minimal representation among T_FR_ cell populations (*15*). This stark dichotomy suggests that T_FR_ cells do not directly engage the foreign antigens driving GC responses. An alternative, though equally plausible, explanation is that strong TCR stimulation by foreign antigens induces T_reg_ cells lineage instability, precluding their identification as conventional T_FR_ cells(*11*).

The failure of T_reg_ cells in aging and autoimmunity presents a critical unresolved question(*16*). With advancing age, progressive erosion of immune homeostasis correlates with increased autoimmune susceptibility, with mounting evidence implicating immunoglobulin-mediated senescence pathways(*17*). Particularly noteworthy are foreign antigens derived from reactivated endogenous retroviruses (ERVs), which elicit persistent antibody responses in aging tissues and may exacerbate immune dysregulation(*18*). These observations raise an important mechanistic question: whether and how antigenic challenge from pathogens or latent viral reservoirs erodes Treg stability, ultimately leading to the breakdown of immune tolerance.

Our study provides definitive mechanistic insight into how TCR affinity dictates T_reg_-mediated self/non-self-discrimination. Through characterization of an endogenous T_FR_ cells-derived TCR, we establish that high-affinity foreign—but not self—antigens selectively induce Foxp3 instability. Remarkably, aged mice exhibit widespread T_reg_ reprogramming into T_FH_/T_H_1-like effector cells, recapitulating key features of human autoimmune pathogenesis. These findings necessitate a paradigm shift in understanding immune tolerance: T_reg_ cells function not as passive suppressors but as dynamic immunological sentinels, where controlled instability represents an adaptive mechanism to selectively restrain self-reactivity while permitting appropriate foreign responses.

## RESULTS

### High-affinity TCR stimulation promotes tT_reg_ instability

We proposed a signal strength model to explain how T_reg_ cells’ fate is shaped upon encountering diverse antigens in the periphery. To test this hypothesis, we employed a fate-mapping system based on*Foxp3*^ΔCNS1-Cre-Thy1.1^*Rosa26*^loxP-stop-loxP-YFP^ bacterial artificial chromosome (BAC) transgenic mice. In this model the Cre-2A-Thy1.1 fusion protein is driven by Foxp3 regulatory elements lacking the conserved noncoding sequence 1 (CNS1)—an enhancer essential for the peripherally induced Treg (pTreg), but not tT_reg_ cells, generation. Consequently, Cre activity induces recombination at the *Rosa26*^loxP-stop-loxP-YFP^ reporter locus, resulting in permanent YFP expression in all tTregs and their progeny, thereby marking their thymic origin even if Foxp3 expression is later lost.(*11*). To simultaneously monitor current Foxp3 expression,, we crossed *Foxp3*^ΔCNS1-Cre-Thy1.1^*Rosa26*^loxP-stop-loxP-YFP^ mice with *Foxp3*^IRES-RFP^ reporter mice (hereinafter referred to as CNS1-YFP-RFP mice). This dual YFP/RFP configuration allows clear discrimination between stable tTregs (YFP⁺RFP⁺) and ex-Tregs that have lost Foxp3 (YFP⁺RFP⁻), enabling precise analysis of tTreg lineage stability and plasticity during peripheral antigen encounter(Fig. 1A).

**Fig. 1.**
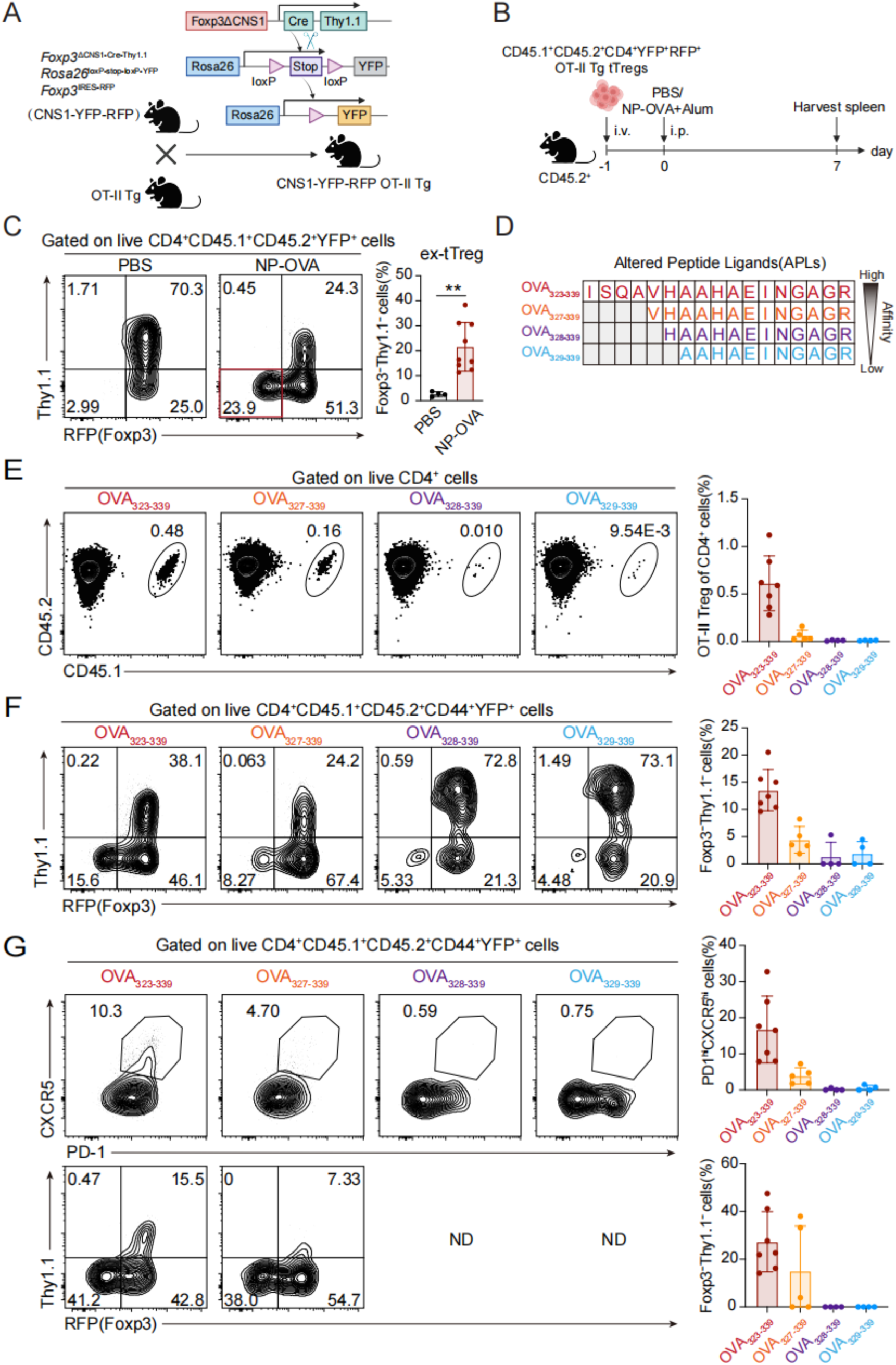
High-affinity TCR stimulation promotes tT_reg_ instability. (**A**) Breeding scheme for generating CNS1-YFP-RFP OT-II Tg mice. *Foxp3*^ΔCNS1-Cre-Thy1.1^ (expressing Cre under the Foxp3 promoter with CNS1 deleted) were crossed with *Rosa26*^loxP-stop-loxP-YFP^ and *Foxp3*^IRES-RFP^ mice. Cre-mediated excision of the loxP-flanked stop cassette at the Rosa26 locus enables YFP expression, while RFP reports Foxp3 expression. CNS1-YFP-RFP mice were further crossed with OT-II TCR transgenic mice (specific for OVA_323–339_). (**B**) Experimental setup for adoptive transfer and immunization. CD45.1^+^CD45.2^+^CD4^+^ YFP^+^RFP^+^ OT-II Tg tT_reg_ cells were transferred intravenously into CD45.2^+^ recipients. One day later, mice were immunized intraperitoneally with PBS or NP-OVA plus alum. Spleens were analyzed by flow cytometry on day 7. (**C**) TCR stimulation influences tT_reg_ cells stability. (left) Flow cytometry plots showing Thy1.1 and Foxp3 expression in CD4^+^YFP^+^ tT_reg_ cells from PBS (n=4) or NP-OVA plus alum (n=9) immunized recipient spleens. Numbers indicate gate frequencies. (right) Quantification of ex-T_reg_ cells (Foxp3^−^Thy1.1^−^, red box). (**D**) Altered peptide ligands (APLs) with varying TCR affinities. OVA-derived APLs are shown with OT-II TCR-binding regions highlighted: OVA_323–339_ (red), OVA_327–339_ (orange), OVA_328–339_ (purple), and OVA_329–339_ (blue). Affinity decreases sequentially. (**E**) Frequency of transferred OT-II Tg T_reg_ cells in recipient mice. Proportion of CD45.1^+^CD45.2^+^ OT-II Tg T_reg_ cells among total CD4^+^ cells after immunization with indicated OVA peptides plus alum. (**F**) Ex-tT_reg_ cells generation correlates with TCR signal strength. Frequency of Foxp3^−^Thy1.1^−^ ex-tT_reg_ cells among activated CD4^+^CD44^+^YFP^+^ cells after peptide immunization. (**G**) Intense TCR signals drive T_FR_ cells instability. Representative flow cytometry plots showing T_FR_ cells differentiation and ex-tT_reg_ cells generation following high-affinity TCR stimulation. ND means not displayed due to being below the detection limit. (E) to (G) show pooled data from two independent experiments with OVA_323–339_ (n=7), OVA_327–339_ (n=5), OVA_328–339_ (n=4), and OVA_329–339_ (n=4). All flow cytometry data were acquired from two independent experiments. Error bars show mean ± SD. *P* values were calculated using the unpaired Student’s *t*-test. \*\**P*<0.01.

To investigate the dynamics of tT_reg_ cells fate, we isolated highly purified tT_reg_ cells (YFP^+^RFP^+^) from OVA-specific OT-II TCR transgenic × CNS1-YFP-RFP mice using fluorescence-activated cell sorting (FACS) and transferred these OT-II TCR-expressing tT_reg_ cells into congenically marked wild-type mice. Recipient mice were immunized intraperitoneally (i.p.) with 4-hydroxy-3-nitrophenyl acetate(NP)-ovalbumin(OVA) precipitated in potassium aluminum sulfate (alum) adjuvant one day after and splenocytes were analyzed after one week (Fig. 1B). NP-OVA stimulation induced a dynamic loss of Thy1.1 and Foxp3 expression in OT-II-expressing tT_reg_ cells compared to phosphate-buffered saline(PBS)-treated controls (Fig. 1C). Notably, Foxp3-negative cells were exclusively observed within the Thy1.1-negative population, suggesting that Thy1.1-negative cells likely represent precursors of ex-T_reg_ cells. These findings demonstrate that TCR stimulation by foreign antigens can activate T_reg_ cells but may also promote their conversion into ex-T_reg_ cells.

To further explore the relationship between signal strength and T_reg_ stability, we employed a series of altered peptide ligands (APLs) derived from OVA (*19*), which exhibit varying affinities for CD4^+^ OT-II Tg T cells (Fig. 1D). Immunization with the highest-affinity OVA_323-339_ peptide resulted in the most significant expansion of OT-II Tg tT_reg_ cells but also the tremendous instability in Foxp3 expression, with the degree of instability closely correlating with TCR-peptide/MHCII affinity (Fig. 1, E and F). Moreover, intense OVA stimulation induced robust differentiation into T_FR_ cells, with ex-T_reg_ cells predominantly localized within the CXCR5^hi^PD-1^hi^ subset (Fig. 1G). These findings support a signal strength model in which high-affinity antigen stimulation activates tT_reg_ cells but also drives their instability and conversion into ex-T_reg_ cells.

### Thymus-derived T_reg_ cells differentiate into T_FR_ cells following foreign antigen stimulation

To further investigate whether endogenous tT_reg_ cells. can recognize foreign antigens and differentiate into T_FR_ cells or ex-T_reg_ cells, we utilized the same tT_reg_ fate-mapping mouse model (CNS1-YFP-RFP). Mice were immunized via footpad injection with NP-OVA precipitated in alum adjuvant to minimize contamination from T_FR_ cells induced by spontaneous GC formation, which can occur due to recognizing follicular B cells-derived autoantigens such as FcγRIIB(*20*). The differentiation of T_FR_ cells in draining and non-draining popliteal lymph nodes was assessed by flow cytometry at 4-, 7-, 10-, 14-, and 21-days post-immunization (Fig. 2A). We found that the percentage and number of YFP-labeled T_FR_ cells significantly increased in the draining popliteal lymph nodes(dpoLN), peaking at day 10 post-immunization and then gradually declining (Fig. 2B), a pattern similar to that observed in T_FH_ cells (fig. S1A). In contrast, very few T_FR_ or T_FH_ cells can be detected in the non-draining popliteal lymph nodes(ndpoLN).

**Fig. 2.**
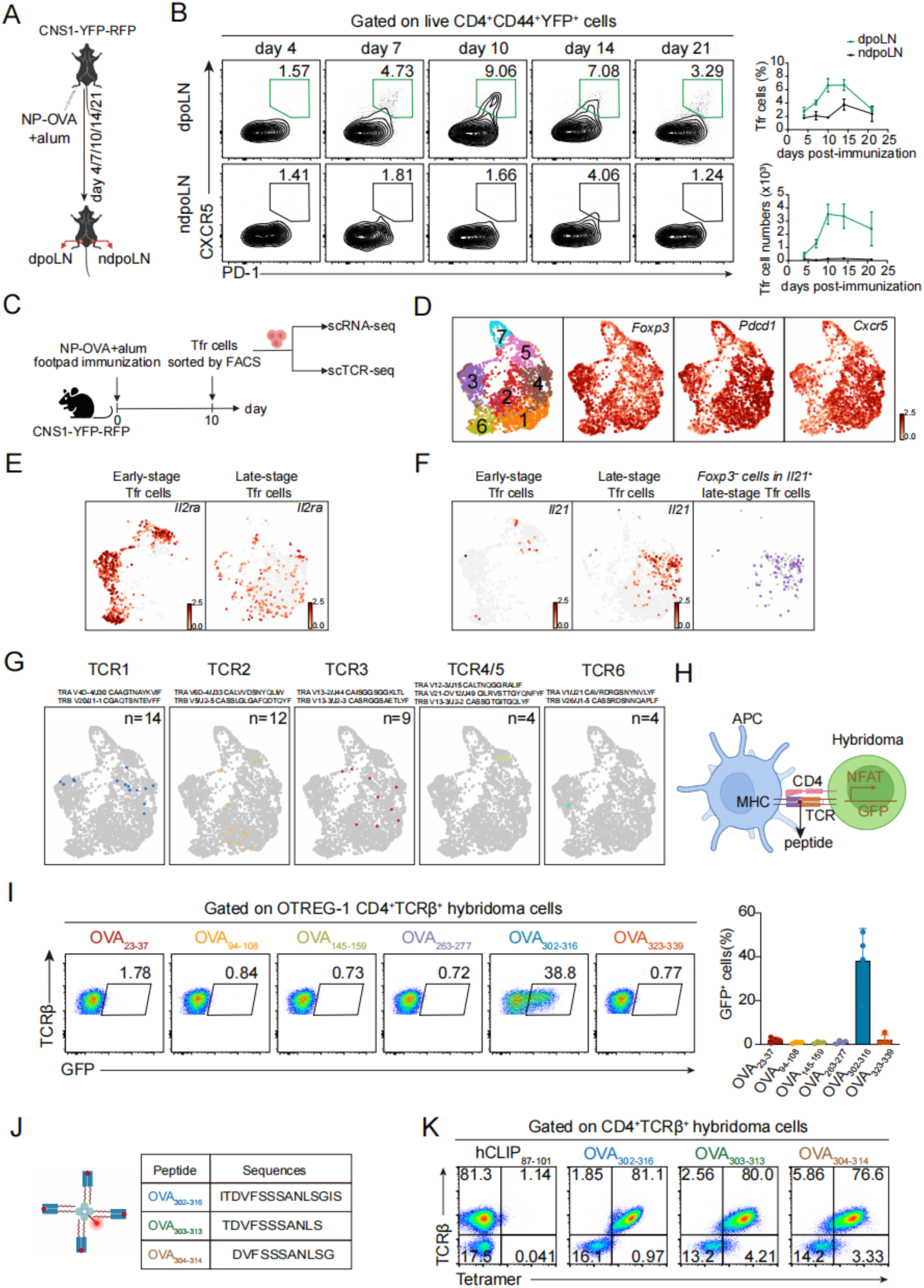
Foreign antigen stimulation drives tT_reg_ differentiation into T_FR_ cells. (**A**) Immunization and analysis scheme. CNS1-YFP-RFP mice received footpad injections of 10 μg NP-OVA plus alum. Draining popliteal lymph nodes (dpoLNs) and non-draining popliteal lymph nodes(ndpoLNs) were analyzed by flow cytometry for PD-1 and CXCR5 expression on CD4^+^YFP^+^ tT_reg_ cells at indicated time points. (**B**) T_FR_ differentiation kinetics. (left) Representative flow plots of PD-1 versus CXCR5 on CD4^+^CD44^+^YFP^+^ tT_reg_ cells. (right) Quantification of T_FR_ cell frequency (top) and numbers (bottom) in dpoLNs versus ndpoLNs. Day4 (n=3), day7 (n=5), day10 (n=4), day14 (n=3), day21(n=2). Flow cytometry data were acquired from two independent experiments. (**C**) scRNA-seq experimental design. PD-1^+^CXCR5^+^YFP^+^ T_FR_ cells from immunized mice were FACS-sorted for single-cell RNA and TCR sequencing. (**D**) T_FR_ cell subsets heterogeneity. UMAP projection of 7 transcriptionally distinct clusters. Dot plot shows relative expression of *Foxp3, Pdcd1* (PD-1), and *Cxcr5* (red: high; gray: low). (**E**) Developmental regulation of CD25. *Il2ra* (CD25) expression patterns in early-(C3/C5/C6) versus late-stage (C1/C2/C4) T_FR_ cell clusters. (**F**) Ex-T_reg_ cytokine production. *Il21* expression map showing Foxp3^-^Il21^+^ cells (purple) within late-stage T_FR_ cell clusters. (**G**) TCR clonal analysis. Six expanded clonotypes (colored) with corresponding CDR3α/β sequences. (**H**) TCR signaling assay. Schematic of NFAT-GFP reporter system in TCR-transduced hybridomas cultured with APCs plus antigen/LPS. (**I**) Antigen specificity validation. GFP induction in OTREG-1 hybridomas stimulated with OVA peptide variants. Representative flow cytometry data of four independent experiment(left). Each data point represents one independent experiment. (**J**) Tetramer reagents. Designed OVA peptide sequences (OVA_302-316_, OVA_303-313_, OVA_304-314_) for TCR specificity testing. (**K**) Tetramer binding confirmation. Flow cytometry analysis of TCRβ versus OVA-tetramer staining (control: hCLIP_87-101_ tetramer).

Subsequently, we aim to determine the TCR specificity for those expanded T_FR_ cells. To overcome this restriction of tetramer-based analysis, we performed single-cell RNA sequencing and paired TCR sequencing on CD4^+^YFP^+^CD44^+^PD-1^hi^CXCR5^hi^ T_FR_ cells isolated from dpoLN at day 10 post-immunization using the 10X Genomics platform (Fig. 2C). We obtained high-quality data from 3,240 T_FR_ cells after removing low-quality cells. Despite three key T_FR_ cell markers—*Foxp3*, *Pdcd1*, and *Cxcr5*—being highly expressed in T_FR_ cells (*7, 8*), these cells did exhibit unpredicted heterogeneity (Fig. 2D). Seven cell clusters were identified and visualized using Uniform Manifold Approximation and Projection (UMAP), a nonlinear dimensionality-reduction technique that groups similar cell populations (fig. S1B). Cluster C7 represented highly proliferative cells expressing high levels of *Top2a* and *Mki67* (fig. S1B). Apart from these highly proliferative cells, T_FR_ cells were divided into two major groups: early-stage CD25^hi^ T_FR_ cells and late-stage CD25^lo^ T_FR_ cells (Fig. 2E), consistent with previous findings (*21*). Early-stage CD25^hi^ T_FR_ cells comprised three significant clusters (C3, C5 and C6). Cells in C3 expressed high levels of *Ccr2*, *Itgae* (encoding CD103), and *Icos*. Cells in C5 expressed high levels of *Il1r2*, while cells in C6 expressed high levels of *Ccr7* and *Sell* (encoding CD62L). Late-stage CD25^lo^ T_FR_ cells also comprised three subsets (C1, C2 and C4). Cells in C1 expressed high levels of *Tcf7* (encoding TCF1), *Tox2*, and *Sh2d1a*, while cells in C2 expressed high levels of the oxysterol receptor *Gpr183* (also known as EBI2), which facilitates T cell positioning to germinal centers, and cells in C4 expressed the highest levels of *Pdcd1*, *Cxcr5*, and *Bcl6* (Fig. 2D and fig. S1B). The differentially expressed genes (DEGs) between CD25^lo^ T_FR_ cells and CD25^hi^ T_FR_ cells overlapped with those between clusters C1, C2, C4 and C3, C5, C6, *Tox2*, *Dennd2d*, *Cxcr5*, *Rnf19a*, *Tnfsf11* and *Asap1* were highly expressed in late-stage T_FR_ cells, while *Il2ra*, *Ccr8*, *Capg*, *Tnfrsf9* and *Vim* poorly expressed in late-stage T_FR_ cells, indicating they are germinal center (GC)-T_FR_ cells (fig. S1C). Interestingly, *Il21*-expressing cells were also identified among late-stage CD25^lo^ T_FR_ cells but few in CD25^hi^ T_FR_ cells, some of which had lost Foxp3 expression, indicating ex-T_reg_ cells are indeed generated in this setting (Fig. 2F).

Next, we focused on the TCR repertoire of T_FR_ cells. Many TCR clones were expanded (detected more than twice). Six expanded TCR clonotypes with distinct transcriptional profiles were synthesized to test the specificity of these expanded TCR clones following NP-OVA immunization (Fig. 2G). The TCRα and TCRβ pairs were linked by a self-cleaving 2A peptide, enabling efficient co-translational cleavage, and cloned into a retroviral vector, named as murine stem cell virus (MSCV)-internal ribosomal entry site (IRES)-nerve growth factor receptor (NGFR). These six TCRα and TCRβ pairs were separately expressed in a TCR-deficient CD4^+^ hybridoma cell line (58α^-^β^-^) to identify antigen-specific TCRs. After co-culturing with or without antigen-presenting cells and NP-OVA for 48 hours, NF-AT response, indicated by GFP induction in TCR-expressing hybridoma cells, was analyzed by flow cytometry (Fig. 2H). Notably, although all TCRs were expressed, only hybridoma cells expressing TCR1 (TRAV4-TRBV20), hereafter named as OTREG-1, induced GFP expression, indicating specificity for OVA (fig. S1D). Notably, OTREG-1(OVA-specific Treg TCR1) was the top expanded clone identified from scRNA-seq, found in both early-stage CD25^hi^ and late-stage CD25^lo^ T_FR_ groups, and some are even IL-21^+^(Fig. 2G and fig. S1E). To further identify the specific epitope recognized by OTREG-1, we designed six I-A^b^-restricted OVA peptides, each 15 amino acids long, using the NetMHCIIpan-4.0 server (fig. S2A). These peptides displayed varying predicted affinities for MHC class II. The classical OT-II-specific OVA_323-339_ peptide was included as a control. TCR activation assays revealed that OTREG-1 specifically recognized the OVA_302-316_ peptide but not others (Fig. 2I). Additionally, we found several tetramers showed the robust binding capacity of OTREG-1 hybridoma cells, including a full-length OVA_302-316_ peptide-specific tetramer and two internal peptides-specific tetramers (OVA_303-313_ and OVA_304-314_), but not the negative control hCLIP-specific tetramer (CLIP_87-101_) (Fig. 2, J and K).

Furthermore, we found that OTREG-1 has a superior antigen sensitivity. To compare OTREG-1 derived from T_FR_ cells with OT-II derived from T_conv_ cells, we generated hybridoma cells expressing OT-II TCR. The results showed that OTREG-1 and OT-II specifically recognized OVA_302-316_ and OVA_323-339_, respectively, without cross-reactivity (fig. S2, B and C). OTREG-1 was easily activated even at 0.1 ug/mL NP-OVA and OVA_302-316_, a very low antigen concentration for T cell activation (fig. S2, D and E). Together, these findings reveal that thymus-derived regulatory T cells possess the capacity to recognize foreign antigens and, upon activation, can undergo clonal expansion and progressive differentiation into germinal-center-associated TFR and ex-Treg..

### OVA_302-316_-specific OTREG-1 exhibits both self and non-self-reactivity

Next, we investigated whether this foreign antigen reactive TCR, OTREG-1, would likewise have self-reactivity since it was cloned from tT_reg_ cells, which are considered to develop following high affinity self-antigen recognition. Thus, we generated an OTREG-1 transgenic (OTREG-1 Tg) mouse carrying the entire TCRα and TCRβ sequences on a C57BL/6J background (fig. S3A). A positive founder was screened and confirmed by specific primers targeting the CDR3 regions and constant regions of TCRα and TCRβ (fig. S3B). Interestingly, when male OTREG-1 Tg mice were mated with female C57BL/6J mice, all male offspring were wildtype mice, while all female offspring were OTREG-1 Tg mice(fig. S3C). According to Mendel’s inheritance laws, this pattern occurs only when the transgenic OTREG-1 is inserted on the X chromosome (fig. S3D).

We next examined the development of OTREG-1-expressing cells in the thymus in male and female OTREG-1 Tg mice, in which approximately 50% of cells express OTREG-1 TCR due to random X chromosome inactivation (fig S3E). Interestingly, although the percentage of CD4^+^CD8^+^ double positive (DP) cells increased in male OTREG-1 Tg mice, the percentage of CD4^+^ single positive (SP) cells sharply decreased (Fig. 3A), which is quite unusual because CD4^+^ T cell-derived TCR transgenic mice typically show significant CD4 lineage bias. We reasoned that OTREG-1-expressing CD4^+^ SP cells might undergo clonal deletion after recognizing self-antigens in the thymus. Thus, we measured the expression of CD24 and Qa2 in TCRβ^hi^ CD4^+^ SP cells, which indicates cell maturity during the thymic SP stage(*22*). We found that the percentage of mature CD24^lo^Qa2^hi^ CD4^+^ thymocytes was decreased dose-dependent (Fig. 3B), indicating that OTREG-1-expressing CD4^+^ cells recognize unknown self-antigens and undertake negative selection. These data support the notion that tT_reg_-derived TCR OTREG-1 possesses dual reactivity, recognizing both self and foreign antigens.

**Fig. 3.**
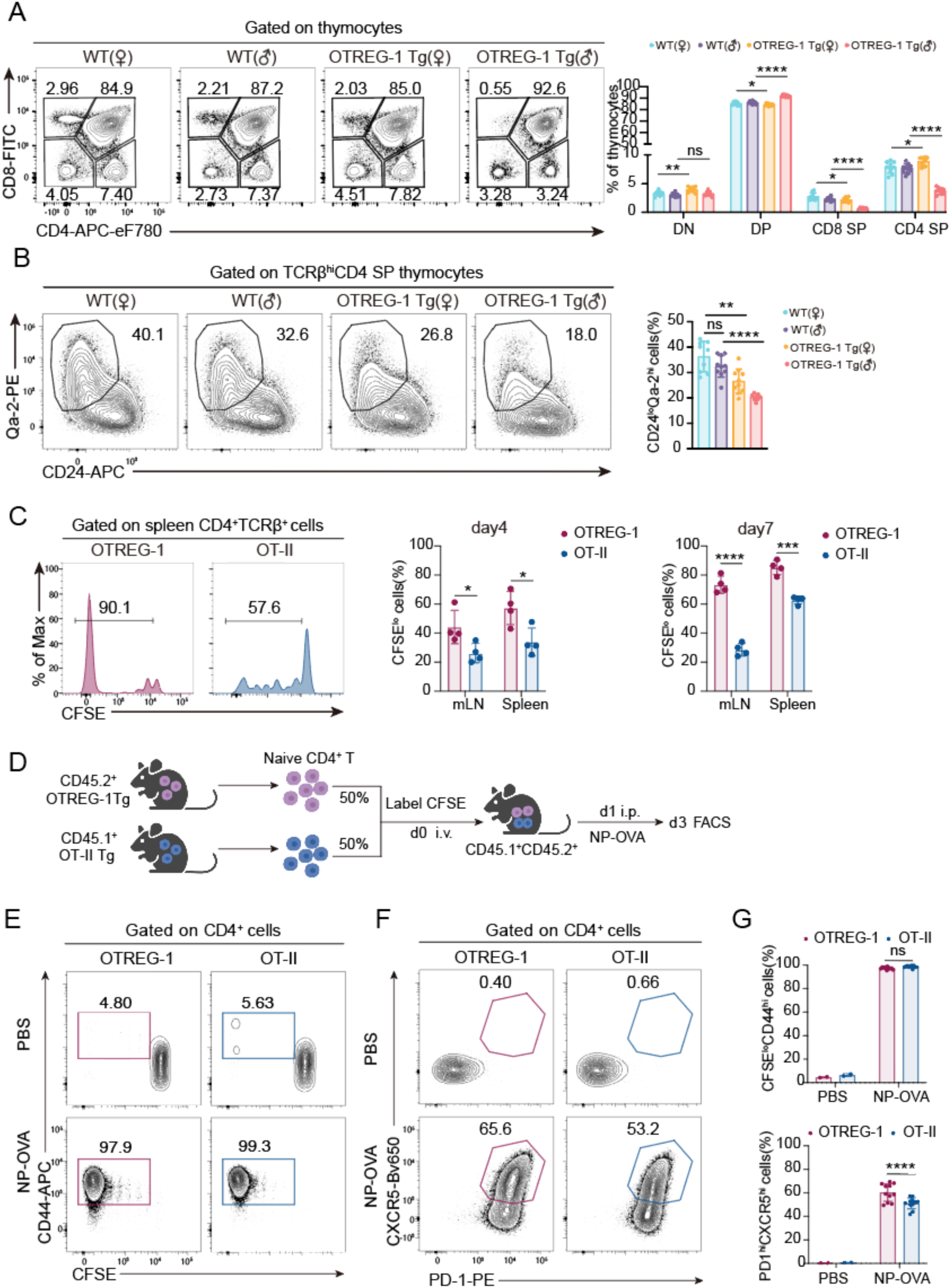
OVA_302-316_-specific OTREG-1 TCR exhibits dual self/non-self reactivity. (**A**) Altered thymocyte development in OTREG-1 Tg mice. (left) Flow cytometry plots of CD4 and CD8 expression in thymocytes from 4–5-week-old WT and OTREG-1 Tg mice. (right) Quantification of DN (double-negative), DP (double-positive), CD4 SP (single-positive), and CD8 SP populations. Data are pooled from all mice. Female WT(n=9), Male WT(n=10), Female OTREG-1 Tg (n=10), Male OTREG-1 Tg (n=10). (**B**) Impaired thymic maturation of OTREG-1 Tg T cells. (left) CD24 and Qa-2 expression on TCRβ^hi^ CD4 SP thymocytes. (right) Frequency of mature CD24^lo^Qa-2^hi^ cells among TCRβ^hi^ CD4 SP thymocytes. Female WT(n=9), Male WT(n=10), Female OTREG-1 Tg (n=10), Male OTREG-1 Tg (n=10). (**C**) Homeostatic proliferation of OTREG-1 Tg T cells. (left) Histograms showing proliferation (CFSE dilution) of splenic OTREG-1 Tg (n=4) and OT-II Tg (n=4) CD4^+^ T cells. (right) Quantification of divided cells at day 4 and day 7. (**D**) Competitive proliferation assay design. Naive CD4^+^CD25^−^CD44^lo^CD62^hi^ T cells from CD45.2^+^ OTREG-1 Tg and CD45.1^+^ OT-II Tg mice were mixed 1:1, CFSE-labeled, and transferred into CD45.1^+^CD45.2^+^ WT hosts. Recipients were immunized i.p. with PBS (n=2) or NP-OVA plus alum (n=11); donor cells were analyzed 3 days later. (**E**) Activation of OTREG-1 versus OT-II Tg T cells. Flow cytometry plots of CFSE dilution (proliferation) and CD44 expression in donor CD45.2^+^ OTREG-1 and CD45.1^+^ OT-II Tg CD4^+^ T cells. (**F**) T_FH_ differentiation potential. Flow cytometry plots of PD-1 and CXCR5 expression on donor OTREG-1 and OT-II Tg CD4^+^ T cells. (**G**) Quantification of activated/proliferating T cells. (upper) Frequency of CFSE^lo^CD44^hi^ cells among OTREG-1 (CD45.2^+^) and OT-II (CD45.1^+^) Tg CD4^+^ T cells. (lower) Frequency of PD-1^hi^CXCR5^hi^ cells among OTREG-1 (CD45.2^+^) and OT-II (CD45.1^+^) Tg CD4^+^ T cells. All flow cytometry data were acquired from at least two independent experiments. Error bars show mean ± SD, *P* values were calculated using unpaired Student’s *t*-test, not significant(ns), \**P*<0.05, \*\**P*<0.01, \*\*\**P*<0.001, \*\*\*\**P*<0.0001.

Our investigation into the behavior of mature CD4^+^ T cells obtained from the lymph nodes and spleen, the secondary lymphoid organ, yielded consistent results. The OTREG-1 Tg mice showed no apparent signs of autoimmunity. However, when we conducted an adoptive transfer of naïve CD4^+^ OTREG-1 cells and control CD4^+^ OT-II cells into syngeneic Rag1^−/−^ mice, we observed that the CD4^+^ OTREG-1 cells exhibited significantly greater proliferation compared to the CD4^+^ OT-II cells after 4 and 7 days in the syngeneic host (Fig. 3C). This finding suggests that, although negative selection has effectively eliminated strongly self-reactive T cells, the OTREG-1-expressing CD4^+^ T cells retain a degree of self-reactivity in the periphery.

As expected, OTREG-1 Tg CD4^+^ T cells can respond with foreign antigen NP-OVA both *in vitro* and *in vivo*. As shown in fig. S3F, CellTrace Violet (CTV)-labeled naïve CD4^+^CD25^-^CD44^lo^CD62L^hi^ OTREG-1 Tg cells were co-cultured with T cells-depleted splenocytes in the presence of OVA_302-316_ or OVA_323-339_ for 3 days. Flow cytometry analysis revealed that CD4^+^OTREG-1 Tg cells were activated (CD44 upregulation) and proliferated (CTV dilution) in response to OVA_302-316_ but not OVA_323-339_ (fig. S3G). Similarly, *in vivo* responses, CFSE-labeled naïve OVA_302-316_-specific CD4^+^OTREG-1 Tg T cells were mixed with OVA_323-339_-specific CD4^+^OT-II Tg T cells at a 1:1 ratio and transferred into wildtype mice, which were immunized the following day (Fig. 3D). Both CD4^+^OTREG-1 Tg and CD4^+^OT-II Tg T cells proliferated and differentiated into T_FH_ cells by day 3 post-immunization (Fig. 3, E to G).

Notably, a higher proportion of CD4^+^OTREG-1-Tg T cells upregulated PD-1 and CXCR5 (Fig. 3, F and G), reflecting their stronger antigen-responsive ability, consistent with their high antigen sensitivity observed in previous hybridoma assays.

Comparable results were obtained using nanoparticle antigen AP205-RBD-OVA_302-339_, in which the 302–339 fragment (including epitopes for OTREG-1 and OT-II) of the OVA was linked to the SARS-CoV-2 spike receptor-binding domain(RBD) and conjugated with acinetobacter phage AP205 using the SpyTag/SpyCatcher system(*23, 24*). Following AP205-RBD-OVA_302-339_ immunization, a higher percentage of CD4^+^OTREG-1 Tg cells upregulated PD-1 and CXCR5 compared to CD4^+^OT-II Tg cells (fig. S3, H and I). Together, these findings indicate that the tTreg-derived TCR OTREG-1 is strongly responsive to foreign antigen while retaining measurable self-reactivity—sufficient to trigger partial negative selection in the thymus but allowing peripheral persistence as a self-tolerant, antigen-sensitive population.

### Distinct responses of OTREG-1 T_reg_ cells to self- versus non-self-antigens

We set out to investigate how the strength of TCR signaling influences the fate of OTREG-1–expressing tTreg cells in response to self and foreign antigens.To address this question, we crossed OTREG-1 transgenic (OTREG-1 Tg) mice with *Foxp3*^ΔCNS1-Cre-Thy1.1^*Rosa26*^loxP-stop-loxP-YFP^ mice to investigate whether OTREG-1-expressing tT_reg_ cells differentiate into ex-T_reg_ cells following foreign antigen stimulation. As illustrated in Fig. 4A, YFP^+^OTREG-1 Tg tT_reg_ cells were adoptively transferred into congenic C57BL/6J mice. Recipient mice were either left unimmunized (to mimic self-antigen engagement) or immunized with NP-OVA precipitated in alum (to simulate foreign antigen exposure) and analyzed after one week.

**Fig. 4.**
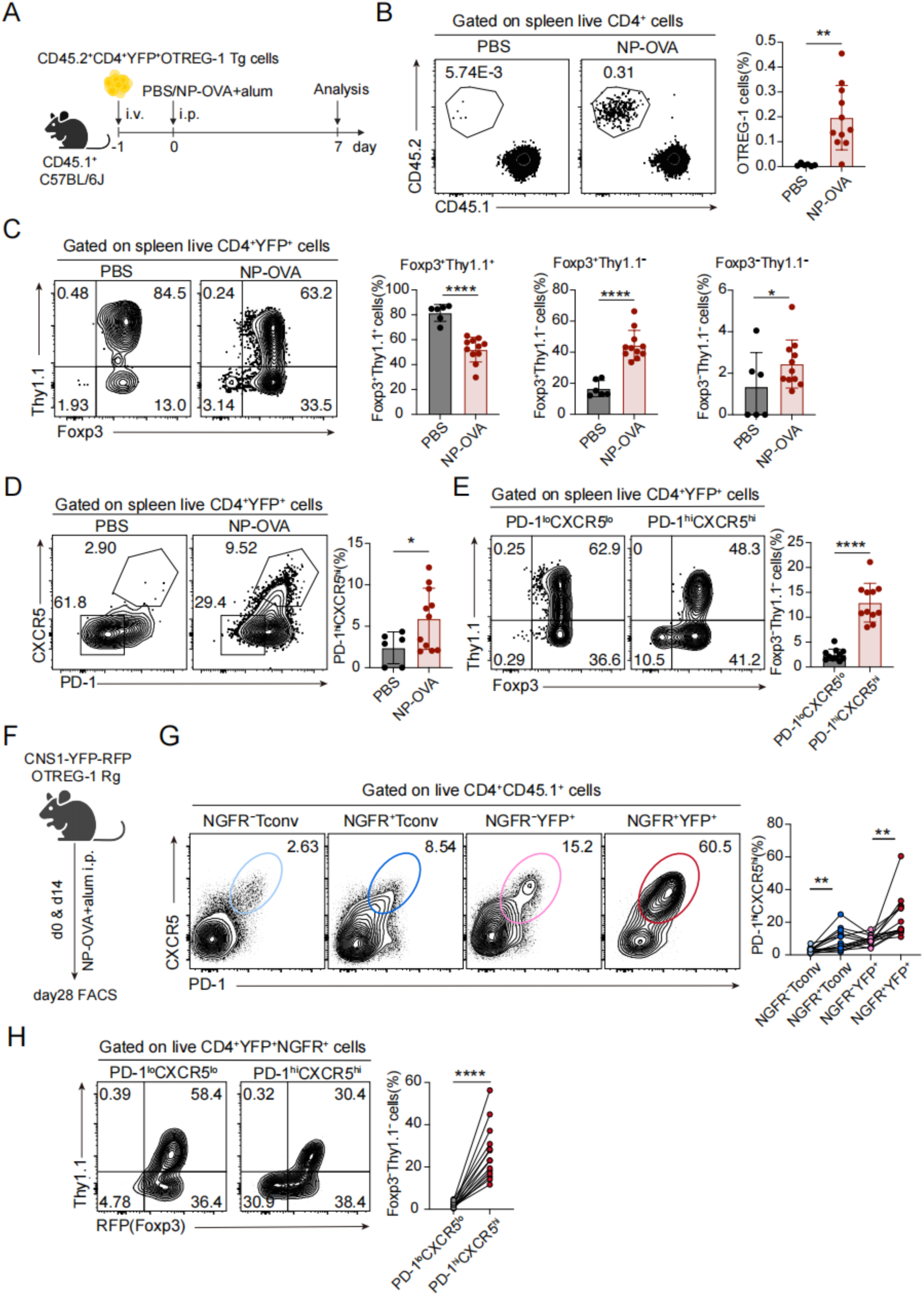
OTREG-1 T_reg_ cells exhibit distinct responses to self-versus foreign antigens. (**A**) Experimental design for assessing the stability of OTREG-1 T_reg_ cells. CD45.2^+^CD4^+^YFP^+^ OTREG-1 Tg tT_reg_ cells were sorted from CNS1-YFP-RFP OTREG-1 Tg mice and adoptively transferred into CD45.1^+^ WT recipients via tail vein injection. Mice were immunized i.p. with NP-OVA plus alum or PBS (control) the following day, and donor cells were analyzed by flow cytometry 7 days post-transfer. (**B**) Homeostasis of transferred tT_reg_ cells. Frequency of donor OTREG-1 Tg tT_reg_ cells (CD45.2^+^) among splenic CD4^+^ T cells in recipient mice. (**C**) Stability of transferred tT_reg_ cells. Foxp3 and Thy1.1 expression in donor tT_reg_ cells from recipient spleens. Numbers indicate gate frequencies. (**D**) T_FR_ cells differentiation. PD-1 and CXCR5 expression on donor tT_reg_ cells following immunization. The percentage of PD-1^hi^CXCR5^hi^ Tfr cells is shown. (**E**) Foxp3 stability of T_FR_ versus non-T_FR_ cell populations. Foxp3 expression in OVA-specific PD-1^lo^CXCR5^lo^ (non-T_FR_) and PD-1^hi^CXCR5^hi^ (T_FR_) cells from immunized mice. (**F**) Antigen challenge protocol for OTREG-1 retrogenic mice. CNS1-YFP-RFP OTREG-1 retrogenic mice received NP-OVA plus alum immunizations at day 0 and day 14, with analysis performed 14 days after the second immunization. (**G**) Antigen-specific versus bystander activation. PD-1 and CXCR5 upregulation on NGFR^+^ CD4^+^ Tconv cells and tT_reg_ cells from retrogenic mice, comparing OVA-specific (NGFR^+^) versus non-specific (NGFR^-^) populations. (**H**) Foxp3 instability in T_FR_ versus non-T_FR_ cell subsets. (left) Flow cytometry analysis of Foxp3 expression in PD-1^lo^CXCR5^lo^ and PD-1^hi^CXCR5^hi^ cell populations gated on CD4^+^CD45.1^+^YFP^+^NGFR^+^ cells. (right) Quantification of ex-tT_reg_ cells (Foxp3^-^Thy1.1^-^) in each subset. (B) to (E) shows pooled data from three independent experiments with PBS group (n=6) and NP-OVA group (n=11). (G) to (H) shows pooled data from three independent experiments (n=14). All flow cytometry data were acquired from at least two independent experiments. Error bars show mean ± SD, *P* values were calculated using paired (G to H) or unpaired (B to E) Student’s *t*-test, \**P*<0.05, \*\**P*<0.01, \*\*\**P*<0.001, \*\*\*\**P*<0.0001.

Compared to unimmunized controls, immunized mice exhibited significant activation and expansion of YFP^+^OTREG-1 Tg tT_reg_ cells, as evidenced by increased cell percentages (Fig. 4B). Notably, YFP^+^ cells in immunized mice showed marked downregulation of Thy1.1-Cre expression, and a substantial proportion of Thy1.1-negative OTREG-1 Tg tT_reg_ cells lost Foxp3 expression in the spleen (Fig. 4C). This finding is consistent with our previous observations that the Thy1.1 marker distinguishes stable T_reg_ cells from those prone to losing regulatory function. Additionally, a significant fraction of YFP^+^OTREG-1 Tg tT_reg_ cells. upregulated PD-1 and CXCR5, indicating differentiation into T_FR_ cells (Fig. 4D). Consistently, T_FR_ cells (CD4^+^YFP^+^PD-1^hi^CXCR5^hi^) were more likely to lose Foxp3 expression compared to non-T_FR_ cells (CD4^+^YFP^+^PD-1^lo^CXCR5^lo^), suggesting that T_FR_ differentiation is associated with increased instability in Foxp3 expression (Fig. 4E).

To eliminate potential artifacts associated with the adoptive transfer, we generated OTREG-1 retrogenic (Rg) mice by infecting bone marrow stem cells from CNS1-YFP-RFP mice with a retroviral vector (MSCV-OTREG-1-IRES-NGFR) and transferring them into irradiated wild-type mice (fig. S4A). We carefully controlled the titer of the virus, allowing only a small percentage of cells to be infected (fig. S4B). After 8–12 weeks of immune reconstitution, Rg mice were immunized with NP-OVA precipitated in alum on days 0 and 14 and analyzed two weeks later (Fig. 4F). Compared to conventional CD4^+^ T cells that do not express OTREG-1 TCR (NGFR^-^), a significantly higher percentage of NGFR^+^ T_conv_ cells (OTREG-1-expressing) upregulated PD-1 and CXCR5, indicating differentiation into T follicular helper (T_FH_) cells. Strikingly, an even greater proportion of NGFR^+^ tT_reg_ cells differentiated into T_FR_ cells, suggesting that tT_reg_ cells in OTREG-1 Rg mice were more prone to T_FR_ differentiation than those in OTREG-1 Tg mice (Fig. 4G). Consistent with previous observation, approximately 30% of antigen-specific tT_reg_ cells lost Foxp3 expression while maintaining CXCR5^hi^PD-1^hi^ phenotypes (Fig. 4H). Together, these findings demonstrate that OTREG-1–expressing tTreg cells mount a robust response to foreign antigen, undergoing activation, expansion, and differentiation into TFR cells accompanied by partial loss of Foxp3 expression. This behavior underscores the dual potential of tTreg cells—maintaining self-tolerance under weak TCR signals while adopting effector-like phenotypes upon strong foreign-antigen stimulation..

### Ex-T_reg_ cells generation from foreign antigen-specific T_FR_ cells

To further investigate whether the generation of ex-T_reg_ cells from foreign antigen-specific T_FR_ cells is a common phenomenon or an exception, we utilized an influenza virus-specific MHC class II tetramer for analysis. CNS1-YFP-RFP mice were immunized with AP205-RBD-NP_311-325_, a particle antigen in which the 311–325 fragment of the influenza virus nucleoprotein (NP) was linked to the spike receptor-binding domain (RBD) of SARS-CoV-2 and conjugated with acinetobacter phage AP205 using the SpyTag/SpyCatcher system(*23, 24*). NP_311-325_-specific CD4^+^ T cells were analyzed by flow cytometry after tetramer (Tet) enrichment on day 7 post-immunization (Fig. 5A).

**Fig. 5.**
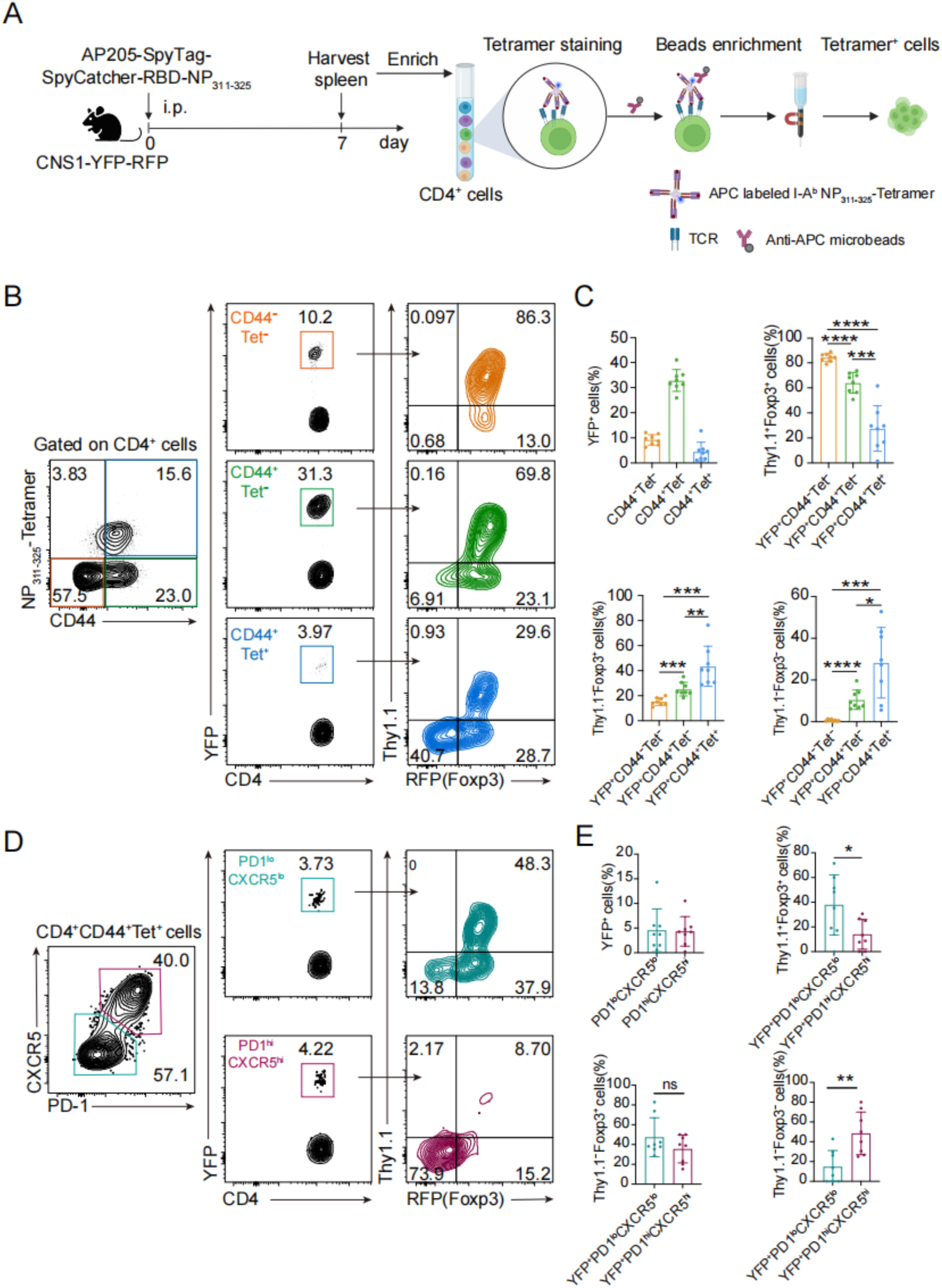
Foreign antigen-specific T_FR_ cells generate ex-T_reg_ cells. (**A**) Experimental design for viral antigen immunization. CNS1-YFP-RFP mice were immunized i.p. with 10 μg AP205-SpyTag-SpyCatcher-RBD-NP_311-325_ virus-like particles. NP_311-325_-specific T cells were subsequently enriched from spleens by magnetic-activated cell sorting (MACS). (**B**) Foxp3 instability in antigen-specific T cells. Flow cytometry analysis of Foxp3 expression in NP_311-325_-tetramer^+^ cells from immunized mice. Influenza A virus PR8-specific T cells were identified using NP_311-325_ tetramer. (**C**) Quantification of **T_reg_** fate decisions. (top left) Frequency of YFP^+^ cells in indicated populations. (top right) Foxp3^+^Thy1.1^+^ stable T_reg_ cells. (bottom left) Foxp3^+^Thy1.1^-^cells. (bottom right) Foxp3^-^Thy1.1^-^ ex-T_reg_ cells in each population. (**D**) Phenotypic analysis of T_FR_ cells stability. PD-1 and CXCR5 expression (left) and corresponding Foxp3 expression (right) in polyclonal NP_311-325_-specific CD4^+^CD44^+^ T cells, showing T_FR_ (PD-1^hi^CXCR5^hi^) versus non-Tfr (PD-1^lo^CXCR5^lo^) cell populations. (**E**) Quantitative assessment of T_reg_ cells fate. (top left) YFP^+^ cell frequency. (top right) Foxp3^+^Thy1.1^+^ cells. (bottom left) Foxp3^+^Thy1.1^-^ cells. (bottom right) Foxp3^-^Thy1.1^-^ex-T_reg_ cells in each population. All flow cytometry data were acquired from at least two independent experiments (n=8). Error bars show mean ± SD, *P* values were calculated using unpaired Student’s *t*-test, not significant(ns), \**P*<0.05, \*\**P*<0.01, \*\*\**P*<0.001, \*\*\*\**P*<0.0001.

Although YFP^+^ tT_reg_ cells were more prevalent among CD44^+^Tet^-^ non-specific cells, YFP^+^ tT_reg_ cells were also detected within the antigen-specific CD44^+^Tet^+^ population. Further analysis revealed that, compared to naïve CD44^-^Tet^-^ tT_reg_ cells and bystander-activated CD44^+^Tet^-^ cells, the percentage of Foxp3^+^Thy1.1^+^YFP^+^ tT_reg_ cells, within the CD44^+^Tet^+^ population substantially decreased. In contrast, the percentage of Foxp3^+^Thy1.1^-^YFP^+^ tT_reg_ cells and ex-T_reg_ cells increased significantly (Fig. 4, B and C). Notably, nearly half of the NP_311-325_-specific CD4^+^ T cells upregulated PD-1 and CXCR5, with approximately 5% being YFP^+^ T_FR_ cells. Consistently, the stable Foxp3^+^Thy1.1^+^ population was strikingly reduced among these cells (Fig. 5, D and E). These results demonstrate that thymus-derived tT_reg_ cells can differentiate into T_FR_ cells following foreign antigen stimulation. However, nearly half of the NP_311-325_-specific YFP^+^ T_FR_ cells lost Foxp3 expression after differentiation, indicating that T_FR_ cells generated in response to foreign antigens are prone to becoming ex-T_reg_ cells.

### tT_reg_ cells exhibit foxp3 instability with aging

Aging is a well-established risk factor for autoimmune disorders, yet its impact on the stability of T_reg_ cells, which are critical for maintaining immune tolerance, remains poorly understood. To investigate this, we analyzed the phenotype of YFP-labeled tT_reg_ cells in young (6–8 weeks old) and aged (8–12 months old) CNS1-YFP-RFP mice under physiological conditions. Significant differences were observed between the two groups: while approximately 75% of tT_reg_ cells in young mice maintained Foxp3 and Thy1.1 expression, aged mice exhibited a remarkable increase in the conversion of tT_reg_ cells into Thy1.1^-^Foxp3^+^ and Thy1.1^-^Foxp3^-^ ex-tT_reg_ cell populations. This instability was particularly pronounced in the spleen, where nearly 10% of tT_reg_ cells lost Foxp3 expression in aged mice. Notably, the ex-tT_reg_ cells induced by aging were exclusively derived from the Thy1.1^-^ cell population, consistent with T_reg_ instability driven by foreign antigen exposure (Fig. 6, A and B). These findings suggest that aging-associated T_reg_ instability contributes to the breakdown of immune tolerance, potentially explaining the increased susceptibility to autoimmune disorders in older individuals.

**Fig. 6.**
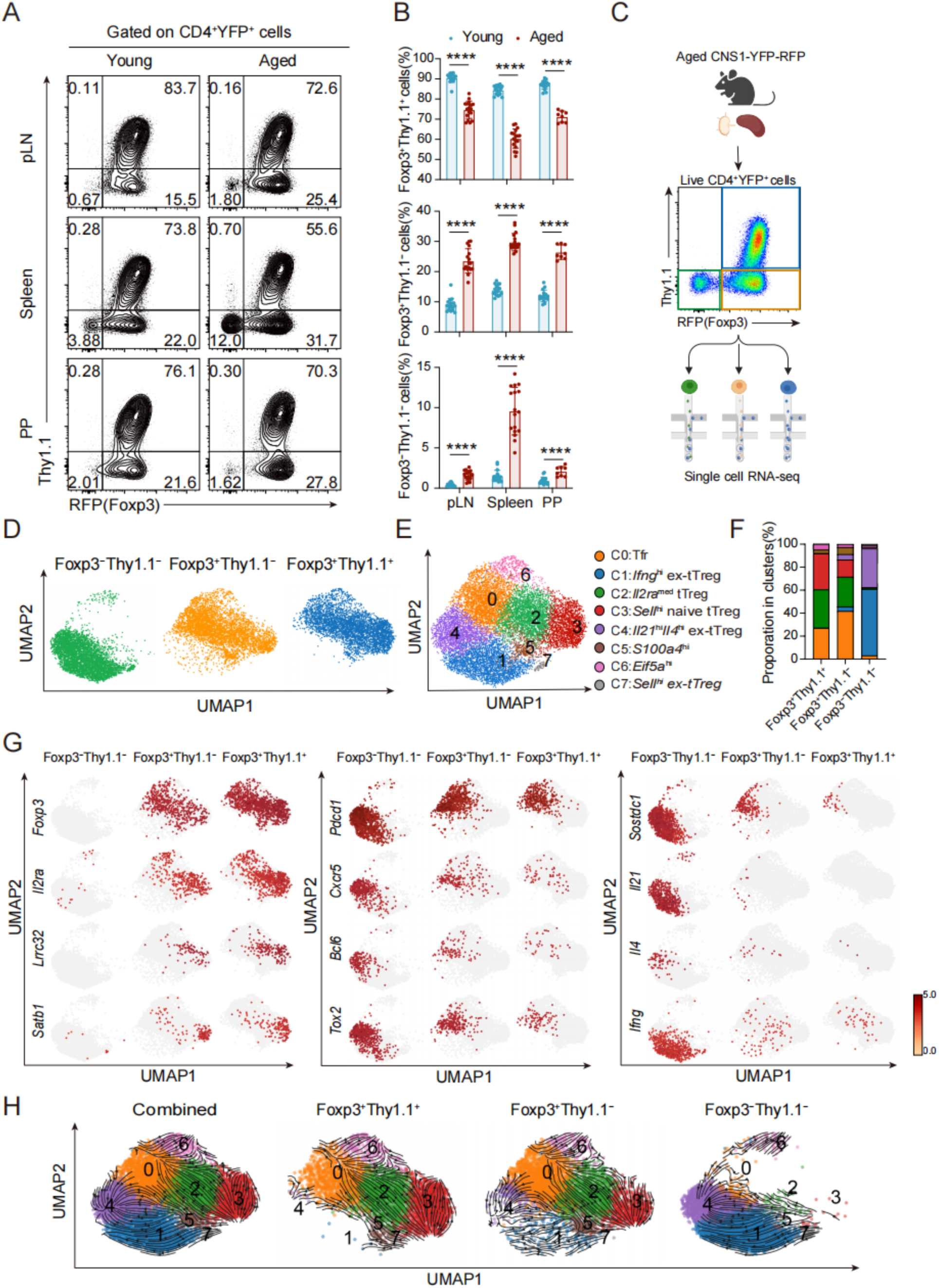
Age-associated Foxp3 instability in tT_reg_ cells. (**A**) Foxp3 stability analysis across lymphoid tissues. Flow cytometry analysis of Foxp3 and Thy1.1 expression in CD4^+^YFP^+^ tT_reg_ cells from peripheral lymph nodes (LNs), spleen, and Peyer’s patches (PP) of young (6-8 week) versus aged (8-12 month) CNS1-YFP-RFP mice. (**B**) Quantification of tT_reg_ fate populations. Percentage of Foxp3^+^Thy1.1^+^ (stable), Foxp3^+^Thy1.1^-^ (unstable), and Foxp3^-^Thy1.1^-^ (ex-tT_reg_) cells among CD4^+^YFP^+^ tT_reg_ cells in indicated tissues, young mice: pLN (n=17), spleen (n=17), PP (n=16), aged mice: pLN (n=17), spleen (n=17), PP (n=7). Data represent mean ± SD. (**C**) Single-cell sorting strategy. Highly purified Foxp3^+^Thy1.1^+^, Foxp3^+^Thy1.1^-^, and Foxp3^-^Thy1.1^-^ (ex-tT_reg_) cells from aged mice were FACS-sorted for scRNA-seq. (**D**) Single-cell transcriptome heterogeneity. UMAP visualization of sorted populations: Foxp3^+^Thy1.1^+^ (blue), Foxp3^+^Thy1.1^-^ (orange), and ex-tT_reg_ cells (green). (**E**) Integrated clustering of tT_reg_ states. UMAP plot showing 8 transcriptionally distinct clusters across all three populations (Foxp3^+^Thy1.1^+^, Foxp3^+^Thy1.1^-^, and ex-tT_reg_ cells). (**F**) Cluster distribution by population. Proportion of cells from each phenotypic population (defined in C) within the 8 transcriptional clusters. (**G**) Key gene expression patterns. UMAP visualization of: Naïve T_reg_ genes (*Foxp3, Il2ra, Lrrc32, Satb1*), effector T_reg_ genes *(Pdcd1, Cxcr5, Bcl6, Tox2*), Conventional CD4^+^ T cell genes (*Sostdc1, Il21, Il4, Ifng*). (**H**) RNA velocity trajectory analysis. Predicted developmental pathways between Foxp3^+^Thy1.1^+^, Foxp3^+^Thy1.1^-^, and ex-tT_reg_ populations based on splicing dynamics. All flow cytometry data were acquired from at least three independent experiments. Error bars show mean ± SD, *P* values were calculated using unpaired Student’s *t*-test, \*\*\*\**P*<0.0001.

To comprehensively characterize the transcriptional profiles and heterogeneity of tT_reg_ subsets, we performed single-cell RNA sequencing (scRNA-seq) on YFP^+^ cells isolated from the spleens of aged mice. Foxp3^+^Thy1.1^+^ cells, Foxp3^+^Thy1.1^-^ cells, and Foxp3^-^Thy1.1^-^ cells (ex-tT_reg_ cells) were sorted from CNS1-YFP-RFP mice and processed using the 10X Genomics system (Fig. 6C). After filtering out low-quality cells, we obtained 13437 high-quality cells (4000 Foxp3^+^Thy1.1^+^ tT_reg_ cells, 4667 Foxp3^+^Thy1.1^-^ tT_reg_ cells, and 4770 ex-tT_reg_ cells). UMAP analysis revealed an almost mutually exclusive distribution between Foxp3^+^Thy1.1^+^ tT_reg_ cells and ex-tT_reg_ cells (Fig. 6D), resembling the segregation between Treg and conventional CD4⁺ T cells described previously, indicating that ex-tTreg cells have fully diverged from the Treg transcriptional program..

As shown in Fig. 6 E-G, Foxp3^+^Thy1.1^+^ tT_reg_ cells were predominantly localized in Foxp3-expressing clusters (C0, C2, C3, C5, and C6), while ex-tT_reg_ cells were concentrated in Foxp3-negative clusters (C1, C4, and C5). Cluster C3 exhibited characteristics of central T_reg_ cells, with high expression of T_reg_ signature genes (e.g., *Foxp3* and *Il2ra*) and the naïve T cell-more activated T_reg_ subsets. Specifically, cells in C0 were *Pdcd1* ^hi^ (encoding PD-1), *Cxcr5*^hi^, and *Bcl6*^+^ (*7, 8*), resembling T_FR_ cells, while cells in C5 expressed high levels of *S100a4 (*data are not shown), marking a novel T_reg_ cell subset. Among the Foxp3-negative clusters, C1 was characterized by high expression of *Ifng* (encoding IFN-γ), a cytokine associated with T_H_1 cells. Cells in C4 expressed *Il21*(encoding IL-21), a T_FH._ cells-associated cytokine(*25*). Gene expression analysis highlighted distinct transcriptional profiles across subsets. Foxp3^+^ tT_reg_ cells expressed T_reg_ cells-specific genes such as *Lrrc32* (encoding GARP) and *Satb1*, particularly in *Sell* ^hi^ (encoding CD62L) C3(*26, 27*). Both ex-tT_reg_ cells and T_FR_ cells expressed T_FH_-associated genes (e.g., *Pdcd1, Cxcr5, Bcl6*, and *Tox2*) and a newly defined T_FH_-associated molecule, *Sostdc1*, which inhibits Wnt-β-catenin signaling to promote T_FR_ differentiation(*28–30*). However, ex-tT_reg_ cells secreted substantial amounts of cytokines associated with T_H_1 cells (e.g., *Ifng*) and T_FH_ cells (e.g., *Il21* and *Il4*) (Fig. 6G).

Foxp3^+^Thy1.1^-^ tT_reg_ cells largely resembled stable Foxp3^+^Thy1.1^+^ tT_reg_ cells, except they had a larger proportion of C0 (T_FR_ cells) and a smaller proportion of C3 (naïve T_reg_ cells) populations. Interestingly, we observed a small subset of Foxp3^+^Thy1.1^-^ tT_reg_ cells exhibiting transcriptional profiles similar to ex-tT_reg_ cells, located in C1 and C4 (Fig. 4D), suggesting these cells may represent precursors of ex-tT_reg_ cells within the T_reg_ cell populations. RNA velocity analysis was used to infer developmental trajectories, revealing that these cells among the Foxp3^+^Thy1.1^-^tT_reg_ cells likely originate from *Pdcd1*^hi^ T_FR_ cell clusters (activated T_reg_ cells) or transition from another subset of activated T_reg_ cells that lose Foxp3 an differentiate into *Il21*^+^*Il4*^+^ex-tT_reg_ cells or *Ifng*^+^ T_H_1-like ex-tT_reg_ cells, respectively (Fig. 4H). These findings demonstrate that aging-induced T_reg_ instability is influenced by T_reg_ activation and specialization.

Supporting this, flow cytometric analysis revealed an age-associated increase inPD-1^hi^CXCR5^hi^ T_FR_ cells (fig S5, A and B). Further analysis revealed that T_FR_ cells were more prone to losing Foxp3 expression, particularly in aged mice (fig S5, A and C). Additionally, aged tT_reg_ cells expressed higher levels of CXCR3(fig S5, D and E), a marker associated with TH1-like Treg cells, and CXCR3⁺ tTregs were more likely to lose Foxp3 expression than CXCR3⁻ counterparts (fig S5, D and F). In summary, our findings reveal that aging promotes Foxp3 instability in tT_reg_ cells, leading to their conversion into ex-tT_reg_ cells with T_H_1- and T_FH_-like phenotypes. This age-associated lineage instability correlates with increased TFR and TH1-Treg populations and may contribute to the gradual breakdown of immune tolerance and heightened susceptibility to autoimmunity in the elderly.

## DISCUSSION

This study defines the mechanisms controlling T_reg_ stability and function, revealing a critical dependence on TCR signal strength, antigen specificity, and aging. We demonstrate that while self-antigen engagement sustains T_reg_ stability, stimulation by non-self-agonists drives instability—a dichotomy mirroring thymic selection principles. The discovery of OTREG-1, a TCR with dual specificity, underscores the importance of intermediate-affinity self-recognition in preserving T_reg_ function. Strikingly, aging exacerbates T_reg_ cells instability, correlating with cumulative foreign antigens exposure. These findings establish a signal strength-dependent model of T_reg_ regulation, offering mechanistic insights into autoimmune pathogenesis and aging-associated immune decline.

Like T_conv_ cells, T_reg_ cells exhibit distinct responses to self- and foreign antigens, likely rooted in thymic selection dynamics. Autoreactive thymocytes undergo initial positive selection in the cortex before encountering high-affinity self-ligands in the medulla. Here, agonist signaling bifurcates: strong, immediate signals drive clonal deletion, whereas delayed, intermediate-affinity interactions promote T_reg_ cells differentiation(*31, 32*). This hierarchy ensures that only T_reg_ cells with optimal self-reactivity persist—equipping them to suppress autoimmunity without compromising foreign immunity. Conversely, excessive TCR stimulation (e.g., by pathogens) disrupts Foxp3 expression, destabilizing T_reg_ cells. Thus, thymic selection calibrates T_reg_ cells to respond to self-antigens while avoiding hyperactivation by foreign stimuli.

Strong TCR signals impair T_reg_ cells’ suppressive capacity by rewiring cytokine receptor expression and downstream pathways. Naïve T_reg_ cells rely on CD25 (IL-2Rα)-STAT5 signaling to maintain lineage fidelity, yet upon activation, they downregulate CD25 and upregulate ICOS, switching to PI3K/AKT-driven signaling(*11, 33*). This transition is epitomized by germinal center follicular regulatory T (GC-T_FR_) cells, which exhibit minimal CD25 but high ICOS expression. Since ICOS-PI3K/AKT activation phosphorylates Foxo proteins—key Foxp3 stabilizers—this shift destabilizes T_reg_ cells(*11, 34, 35*). Fine-tuning TCR signal strength is therefore essential to balance T_reg_ activation and functional integrity.

Aging and autoimmunity are inextricably linked, with immunosenescence fostering a permissive environment for immune dysregulation. In aged mice, T_reg_ instability escalates alongside prolonged foreign antigen exposure, driving the emergence of ex-T_reg_ cells with T_H_1-or T_FH_-like phenotypes. This phenotypic drift may underlie the erosion of self-tolerance in aging, predisposing older individuals to autoimmunity. Notably, chronic antigenic stressors (e.g., endogenous retroviruses) could further exacerbate this process, suggesting that aging-associated T_reg_ dysfunction is both a cause and consequence of persistent immune activation.

Our signal strength-centric model unifies disparate observations on T_reg_ biology, explaining how self-tolerance is maintained without compromising pathogen responsiveness. However, key questions remain: How do aging and chronic inflammation modulate TCR signaling. thresholds in T_reg_ cells? Can therapeutic interventions recalibrate signal strength to restore stability? Addressing these gaps will not only refine our understanding of T_reg_ regulation but also illuminate novel strategies to combat autoimmunity and age-related immune decline.

## MATERIALS AND METHODS

### Study design

This study aimed to investigate how the affinity of TCRs to foreign versus self-antigens influences the stability of T_reg_ cells and the functional implications of T_reg_ instability for immune tolerance. We sought to determine whether high-affinity antigen recognition leads to the reprogramming of T_reg_ cells into effector cells and to characterize the transcriptional landscape associated with this process. To achieve these objectives, we employed a multifaceted approach that included novel transgenic models, adoptive transfers, and single-cell technologies. Our genetic and functional analyses utilized CNS1-YFP-RFP Foxp3 fate-mapping mice to track T_reg_ lineage history. These were crossed with TCR transgenic models, such as OT-II and the endogenous T_FR_-derived OTREG-1, to control for antigen specificity. This methodology enabled us to compare Treg responses to high-affinity foreign and self-antigens. We further validated the phenomenon of antigen-driven instability using a vaccine immunization model..

Additionally, we conducted single-cell RNA sequencing (scRNA-seq) on antigen-induced T_FR_ and aged T_reg_ cells to define the transcriptional continuum from stable T_reg_ cells to ex-T_reg_ cells, identifying key effector genes and developmental trajectories. Our primary assessments included monitoring Foxp3 expression stability through fate-mapping and flow cytometry, evaluating T cell phenotypes for T_FH_, T_H_1, and other effector markers using flow cytometry, analyzing transcriptomic signatures, and tracking the spontaneous emergence of ex-T_reg_ cells in aged mice as a model for human aging. Mice were randomly assigned to experimental groups based on their genotype, and each experiment was conducted at least three times to ensure reproducibility. Details regarding the number of biological and technical replicates, along with the relevant statistical analyses, are provided in the figure legends.

### Mice

The *Foxp3*^ΔCNS1-Cre-Thy1.1^ mouse strain has been previously described (*11*). *Rosa26*^loxP-stop-loxP-YFP^ (*36*) and *Foxp3*^IRES-RFP^ (*5*) strains were obtained from the Jackson Laboratory. C57BL/6J, CD45.1 congenic C57BL/6J, and Rag1^−/−^ mice were purchased from GemPharmatech Co., Ltd (Nanjing, China). OT-II transgenic mice were kindly provided by Mo Xu (National Institute of Biological Sciences, Beijing). The OTREG-1 transgenic mouse line was generated in this study. For experiments involving *Foxp3*^ΔCNS1-Cre-Thy1.1^*Rosa26*^loxP-stop-loxP-YFP^*Foxp3*^IRES-RFP^ (hereafter referred to as CNS1-YFP-RFP) mice, young mice (6–8 weeks old) and aged mice (8–12 months old) were used. All other mice were utilized at 6–12 weeks of age unless otherwise specified. Mice were housed under specific pathogen-free (SPF) conditions at the Institute of Microbiology, Chinese Academy of Sciences. The facility maintained a 12-hour light/dark cycle, with an ambient temperature of 20–22 °C and relative humidity of 40–60%. Euthanasia was performed using CO_2_ asphyxiation, followed by confirmation of death via cessation of heartbeat and respiration. All experimental procedures were approved by the Institutional Animal Care and Use Committee of the Institute of Microbiology, Chinese Academy of Sciences (Permit No. APIMCAS2021104) and conducted in accordance with institutional guidelines.

### Cell lines

The Plat-E cell line was maintained in our laboratory. The NFAT-GFP CD4^hi^58α^-^β^-^ hybridoma cell line (*37*) was generously provided by Mo Xu (National Institute of Biological Sciences, Beijing). Plat-E cells were cultured in Dulbecco’s Modified Eagle’s Medium (DMEM; Gibco) supplemented with 10% fetal bovine serum (FBS; VivaCell) and 1% penicillin-streptomycin (Thermo Fisher Scientific). Hybridoma cells were maintained in RPMI 1640 medium (Gibco) containing 10% FBS (VivaCell), 50 µM 2-mercaptoethanol (Thermo Fisher Scientific), 1× non-essential amino acids (NEAA; Thermo Fisher Scientific), and 1% penicillin-streptomycin. All cell lines were incubated at 37°C in a humidified atmosphere with 5% CO_2_ (Thermo Fisher Scientific incubator).

### Plasmids

The TCRα and TCRβ sequences of TCR1(OTREG-1), TCR2, TCR3, TCR4, TCR5, and TCR6—linked via a 2A peptide—were synthesized by GenScript and subsequently subcloned into an MCSV-IRES-NGFR retroviral vector. Full-length TCRα and TCRβ cDNA sequences were amplified and individually cloned into the pCD2-TCRα and p428-TCRβ vectors, respectively, using homologous recombination.

### Antibodies

The following antibodies were used for flow cytometry analysis. PerCP-Cyanine5.5 mouse anti-mouse CD45.1 (A20, eBioscience, 45-0453-82, 1:400), Alexa Fluor 700 mouse anti-mouse CD45.2 (104, eBioscience, 56-0454-82, 1:400), PerCP-Cyanine5.5 rat anti-mouse CD4 (RM4-5, eBioscience, 45-0042-82, 1:400), APC-eFluor 780 rat anti-mouse CD4 (GK1.5, eBioscience, 47-0041-82, 1:300), PE-Cyanine7 rat anti-mouse CD8a (53-6.7, Biolegend, 100722, 1:500), PerCP-Cyanine5.5 Armenian hamster anti-mouse TCRbeta (H57-597, eBioscience, 45-5961-80, 1:400), APC rat anti-mouse CD44 (IM7, eBioscience, 17-0441-83, 1:200), PE-Cyanine7 rat anti-mouse CD44 (IM7, eBioscience, 25-0441-81, 1:200), APC rat anti-mouse CD62L (MEL-14, eBioscience, 17-0621-82, 1:200), eFluor 450 rat anti-mouse CD25 (PC61.5, eBioscience, 48-0251-82, 1:200), eFluor 450 rat anti-mouse Foxp3 (FJK-16s eBioscience, 48-5773-82, 1:100), Brilliant Violet 421 mouse anti-mouse CD90.1 (Thy1.1) (OX-7, Biolegend, 202529, 1:200), PE-Cyanine7 rat anti-mouse CD279 (PD-1) (RMP1-30, Biolegend, 109110, 1:200), Biotin rat anti-mouse CD185 (CXCR5) (L138D7, Biolegend, 145510, 1:100), Biotin Armenian hamster anti-mouse CXCR3 (CXCR3-173, eBioscience, 13-1831-82, 1:200), PE-Cyanine7 Armenian hamster anti-mouse CXCR3 (CXCR3-173, Biolegend, 126516, 1:200), APC rat anti-mouse CD24 (30-F1, BD Pharmingen, 562349, 1:200), Biotin mouse anti-mouse Qa2 (695H1-9-9, Invitrogen, MA5-28660, 1:200), BV650 Streptavidin (BD Horizon, 563855, 1:200), PE Streptavidin (Biolegend, 405203). APC-labeled Tetramer I-A^b^ human CLIP _87-101_, APC-labeled Tetramer I-A^b^ chicken OVA_302-316_, APC-labeled Tetramer I-A^b^ chicken OVA_303-313_, APC-labeled Tetramer I-A^b^ chicken OVA_304-314_ and APC-labeled Tetramer I-A^b^ Influenza A NP_311-325_ were provided by NIH Tetramer Core Facility and were used at a dilution ratio of 1:100.

### Flow cytometry analysis

Mice were euthanized by CO_2_ asphyxiation followed by cervical dislocation. Single-cell suspensions were prepared from the thymus, lymph nodes, and spleen via mechanical disruption in RPMI 1640 medium supplemented with 2% (v/v) FBS. After centrifugation at 2000 rpm for 2 min at 4°C, cell pellets were resuspended in 1 mL of ACK lysis buffer (150 mM NH_4_Cl, 10 mM KHCO_3_, 0.1 mM EDTA) and incubated for 2 min at room temperature (RT) to lyse red blood cells (RBCs). The suspensions were then diluted in 10 mL of RPMI 1640 with 2% FBS, centrifuged again (2000 rpm, 2 min, 4°C), and resuspended in FACS buffer (PBS containing 0.2% BSA). Cells were filtered through a 70 µm cell strainer before staining. For surface staining, cells were first incubated with anti-mouse CD16/CD32 (Fc block; BioXCell) in FACS buffer on ice for 10 min in the dark. Subsequently, they were stained for 30 min at 4°C with a mixture of fluorochrome-conjugated antibodies and Fixable Viability Dye eFluor 506 (eBioscience). Cells were then fixed with 4% paraformaldehyde (PFA; Santa Cruz) for 8 min at RT. For intracellular staining Foxp3 together with YFP, surface-stained cells were pre-fixed with 1% PFA for 8 min at RT, followed by fixation/permeabilization using Fixation/Permeabilization Buffer (Invitrogen) at 4°C for 30 min. Cells were then stained with anti-Foxp3 antibody in Perm/Wash Buffer (BD Biosciences) at 4°C for 30 min. After washing with Perm/Wash Buffer, cells were resuspended in FACS buffer for analysis. Samples were analyzed on an LSR Fortessa flow cytometer (BD Biosciences), and data were processed using FlowJo v10 software (Treestar).

### Naïve CD4^+^ T cell isolation and purification

Single-cell suspensions were prepared from murine spleen and lymph nodes. CD4^+^ T cells were negatively selected using a two-step magnetic bead separation protocol. First, cells were incubated with anti-CD19 (Bio X Cell) and anti-CD8α (Bio X Cell) monoclonal antibodies, followed by depletion with BioMag goat anti-rat IgG beads (QIAGEN) using a magnetic separator (Stemcell Technologies). The enriched CD4^+^ T cell population was then incubated with 10 μg/mL anti-CD16/CD32 (clone 2.4G2) for 10 minutes at 4°C to block Fc receptors and then stained with a cocktail of surface antibodies for 30 minutes at 4°C. Naïve CD4^+^ T cells (CD4^+^CD25^-^CD44^lo^CD62L^hi^) were subsequently sorted using a FACSAria III cell sorter (BD Biosciences) controlled by FACSDiva software (BD Biosciences), achieving >98% purity as verified by post-sort analysis.

### Tetramer enrichment and staining

All MHC class II tetramers were generously provided by the NIH Tetramer Core Facility. To detect influenza A nucleoprotein (NP)-specific tT_reg_ cells, single-cell suspensions from spleens were first subjected to CD19^+^ B cells and CD8^+^ T cells depletion as previously described (*38*). The enriched CD4^+^ T cell population was then incubated with 10 μg/mL anti-CD16/CD32 (clone 2.4G2) for 10 minutes at 4°C to block Fc receptors. For tetramer staining, cells were incubated with APC-conjugated MHC class II tetramers containing the NP_311-325_ epitope (QVYSLIRPNENPAHK) from influenza A virus PR8 strain for 1 hour at room temperature. Following tetramer staining, cells were washed and incubated with 25 μL anti-APC magnetic microbeads for 30 minutes at 4°C. Tetramer-positive cells were then enriched using magnetized LS columns. The APC-labeled, antigen-specific T cells were subsequently stained with surface antibodies. For intracellular Foxp3 staining, cells were pre-fixed with 1% paraformaldehyde (PFA) for 8 minutes at room temperature, then processed using the Foxp3/Transcription Factor Buffer Set according to the manufacturer’s instructions.

### Soluble antigen preparation and immunization

For antigen-alum precipitation, NP-OVA or altered peptide ligands at predetermined concentrations were mixed with 10% potassium aluminum sulfate (AlK(SO₄)₂·12H₂O) in a 1:1 ratio. The pH was adjusted to 7.0 by dropwise addition of 1 M NaOH to induce precipitation. The mixture was washed three times with phosphate-buffered saline (PBS) and resuspended in PBS for immunization. For footpad immunization, mice received 10 μg of alum-precipitated NP-OVA (Biosearch Technologies) as previously described (*39*). Intraperitoneal immunizations were performed with 200 μg of alum-precipitated NP-OVA or modified OVA peptide ligands.

### T cell proliferation and immunization

Naïve CD45.2^+^ OTREG-1-transgenic CD4^+^ T cells (2×10^5^) were labeled with 10 μM CellTrace Violet (CTV) for 10 min at 37°C and co-cultured with T cell-depleted splenocytes (2×10^5^) in U-bottom 96-well plates for 72 hours with OVA_302-316_ or OVA_323-339_ peptide (0.1-10 μg/mL), with proliferation assessed by CTV dilution; for *in vivo* studies, CFSE-labeled OTREG-1 and OT-II transgenic CD4^+^ T cells (1:1 ratio, 2×10^5^ each) were co-transferred into wild-type mice via tail vein injection, followed by intraperitoneal NP-OVA immunization 24 hours later and CFSE dilution analysis after 72 hours. For immunizations, NP-OVA or altered peptide ligands were mixed in 1:1 ratio with 10% potassium aluminum sulfate (AlK(SO₄)₂·12H₂O), pH-adjusted to 7.0 with 1M NaOH, washed 3 times with PBS, and resuspended for administration either 10 μg in footpads or 200 μg intraperitoneally.

### Pathogen-like antigen preparation and immunization

The pathogen-like antigens AP205-RBD-NP_311-325_ and AP205-RBD-OVA_302-339_ were generated as described (*23*) with modifications: AP205-SpyTag was created by fusing SpyTag to the C-terminus of the major AP205 coat protein (pET21 vector), while synthetic sequences encoding SARS-CoV-2 RBD (aa 319-541; YP_009724390.1) and influenza NP_311-325_ (or OVA_302-339_ for the alternative construct) were cloned into pCEP4; SpyCatcher-RBD-NP_311-325_ was generated by fusing the S protein signal peptide and SpyCatcher to RBD’s N-terminus with a C-terminal 12×His-tag, with NP_311-325_ replaced by OVA_302-339_ for the second construct; following protein purification, 100 μg SpyCatcher-RBD-NP_311-325_ or -OVA_302-339_ was mixed with 400 μg AP205-SpyTag in PBS (≥1h incubation on ice) to form the final antigens, of which 10 μg was administered intraperitoneally per mouse.

### Retroviral transfection and hybridoma generation

For retrovirus production, Plat-E cells were transfected with TCR-expressing plasmids using either Lipofectamine^TM^ 2000 (Invitrogen) or calcium phosphate (Boytime) according to manufacturers’ protocols. Viral supernatants were collected 48 hours post-transfection and filtered through 0.45 μm membranes. TCR hybridomas were generated by cloning synthesized TCRα and TCRβ sequences (OTREG-1, TCR2-6) linked via P2A self-cleavage sequences (GenScript Biotech) into the MSCV-IRES-NGFR retroviral vector. Hybridoma cells were transduced via spin infection (1700 rpm, 45 min, 25°C) using viral supernatants supplemented with 10 μg/mL polybrene.

### Hybridoma activation assay

Antigen-presenting cells (APCs) were prepared by depleting CD4^+^ and CD8^+^ T cells from C57BL/6J mouse splenocytes. For activation assays, 1×10^4^ hybridoma cells were co-cultured with 2×10⁵ APCs in U-bottom 96-well plates containing 1 μg/mL LPS and antigen for 48 hours. TCR activation was quantified by flow cytometric analysis of GFP expression.

### Generation of OTREG-1 transgenic mice

The OTREG-1 transgenic mouse line was generated by pronuclear microinjection of C57BL/6J fertilized oocytes with TCRα and TCRβ gene constructs. Full-length TCRα and TCRβ cDNA sequences were first inserted into pCD2-TCRα and p428-TCRβ vectors, respectively, using homologous recombination. The TCRα construct was linearized with KpnI and XbaI restriction enzymes, while the TCRβ construct was digested with NotI to remove the vector backbone. The purified TCRα and TCRβ fragments were co-injected into fertilized oocytes. Founder mice were identified by PCR genotyping using transgene-specific primers targeting the CDR3 regions (TCRα: 313 bp; TCRβ: 517 bp).

### Generation of OTREG-1 retrogenic mice

Young (6-8 week old)CD45.2^+^ CNS1-YFP-RFP donor mice received intravenous 5-fluorouracil (10 mg/mL, 300 μL per 20 g) on day 1 to deplete proliferating cells, followed by Plat-E cell plating (1.2×10⁶ cells/well in 6-well plates) on day 2 evening and calcium phosphate transfection with MSCV-OTREG-1-IRES-NGFR/pCL-Eco plasmids at day 3 (12 h post-plating); at day 4, donor bone marrow was isolated from femurs/tibiae in RPMI 1640 (2% FBS, 1% penicillin-streptomycin, 25 mM HEPES, 10 μg/mL DNase I), lysed with ACK buffer, filtered (40 μm), and plated in DMEM/15% FBS (1×10⁶ cells/mL, 2 mL/well in 24-well plates); viral supernatants collected at days 5-6 (0.45 μm-filtered, 8 μg/mL polybrene) were used for spinoculation (2400 rpm, 25°C, 2 h) of bone marrow cells cultured with mSCF (100 ng/mL), mIL-6 (50 ng/mL), and mIL-3 (20 ng/mL), while recipient C57BL/6J mice received 6 Gy γ-irradiation (Co⁶⁰) before intravenous transfer of 5×10⁵ transduced cells, with experiments conducted 8-12 weeks post-reconstitution.

### Single-cell isolation and sequencing

For NP-OVA immunized mice, we fluorescence-activated cell sorted (FACS) CD4^+^CD44^+^YFP^+^PD-1^hi^CXCR5^hi^ follicular regulatory T (T_FR_) cells from popliteal lymph nodes at day 10 post-immunization. Washed cells were processed using the Chromium Controller (10X Genomics) with Chromium Single Cell 5’ and V(D)J Library Construction Kits, followed by paired-end sequencing (Illumina NovaSeq 6000). For aged tT_reg_ analysis, CD4^+^ T cells were magnetically enriched (MACS) from lymph nodes and spleens of 12-month-old CNS1-YFP-RFP mice. Three populations (CD4^+^YFP^+^RFP^+^Thy1.1^+^, CD4^+^YFP^+^RFP^+^Thy1.1^-^, CD4^+^YFP^+^RFP^-^Thy1.1^-^) were FACS-purified (>99% purity), washed in PBS/0.4% BSA, and viability-confirmed (>93% by AO/PI staining). Single-cell libraries were prepared using the Chromium Single Cell 3’ Kit (10x Genomics) and sequenced (NovaSeq 6000, 150 bp paired-end).

### Single-cell RNA-seq analysis pipeline

Raw sequencing data were aligned to mm10-2020-A reference genome using Cell Ranger (v4.0.0), followed by quality control excluding cells with <200 detected genes or >10% mitochondrial UMIs and genes detected in <3 cells, with doublet removal via Scrublet(v0.2.3); after library-size normalization (scanpy.pp.normalize_total), we selected 2,000 highly variable genes (scanpy.pp.highly_variable_genes), regressed out technical covariates (total counts, mitochondrial content), performed PCA (100 components, svd_solver=’arpack’), constructed k-nearest neighbor graphs, applied Leiden clustering (resolution=0.6-0.7), generated UMAP visualizations, and identified cluster markers (scanpy.tl.rank_genes_groups).

### RNA velocity analysis

RNA velocity analysis was performed on aged tT_reg_ cells using the following workflow. First, spliced and unspliced transcripts were recalculated using velocyto (v0.2.5). Following count normalization, genes detected in >20 cells across both transcript populations were retained. Highly variable genes were selected (scanpy.pp.highly_variable_genes) for PCA dimensionality reduction. RNA velocity moments were then computed (scvelo.pp.moments), and transcriptional dynamics were estimated using stochastic modeling (scvelo.tl.velocity). Velocity vectors were projected onto single-cell manifolds through cosine similarity transformation (scvelo.tl.velocity_graph) and visualized on UMAP embeddings (scvelo.pl.velocity_embedding_stream).

### Statistical analysis

Data were analyzed by using the GraphPad Prism 9 and the mean ± SD were shown for representative results. Continuous data were compared using paried or unpaired Student’s *t*-tests. \**P*<0.05, \*\**P*<0.01, \*\*\**P*<0.001, \*\*\*\**P*<0. 0001.Throughout the study, all experiments were performed independently at least twice.

## Materials availability

The raw single-cell RNA-seq data presented in this paper have been submitted to the Gene Expression Omnibus database under accession numbers GSE297563 and GSE298156.Any additional information required to reanalyze the data reported in this paper in available from the lead contact upon request.

## Acknowledgments

We thank Dr. Xuguang Tai and Sherif Badr for their critical review of the manuscript and Dr. Zhaolin Hua and Zhongmei Zhang for helpful suggestions. We are grateful to the Pathogenic Microbiology and Immunology Public Technology Service Center for their support and to all members of our laboratory for productive discussions. MHC class II tetramers were provided by the NIH Tetramer Core Facility (NIH Contract 75N93020D00005; RRID:SCR_026557). We also thank Dr. Fengchao Wang for generating the OTREG-1 TCR transgenic mouse.

This work was supported by the National Natural Science Foundation of China (U23A20464) and the National Key Research and Development Program of China (2021YFC2300503). The graphical abstract was created with BioRender (https://biorender.com).

## Author contributions

L.L.,J.G., R.H., X.C., W.X., X.C., and Q.J. conducted experiments. F.Z., B.H. and F.Z. provided experimental assistance. L.L. and X.Z. designed the study and wrote the manuscript. X.Z. supervised the project.

## Declaration of generative AI and AI-assisted technologies in the writing process

During the preparation of this work the author(s) used DeepSeek in order to improve the readability and language of the manuscript. After using this tool/service, the author(s) reviewed and edited the content as needed and take(s) full responsibility for the content of the published article.

---

**fig. S1.**
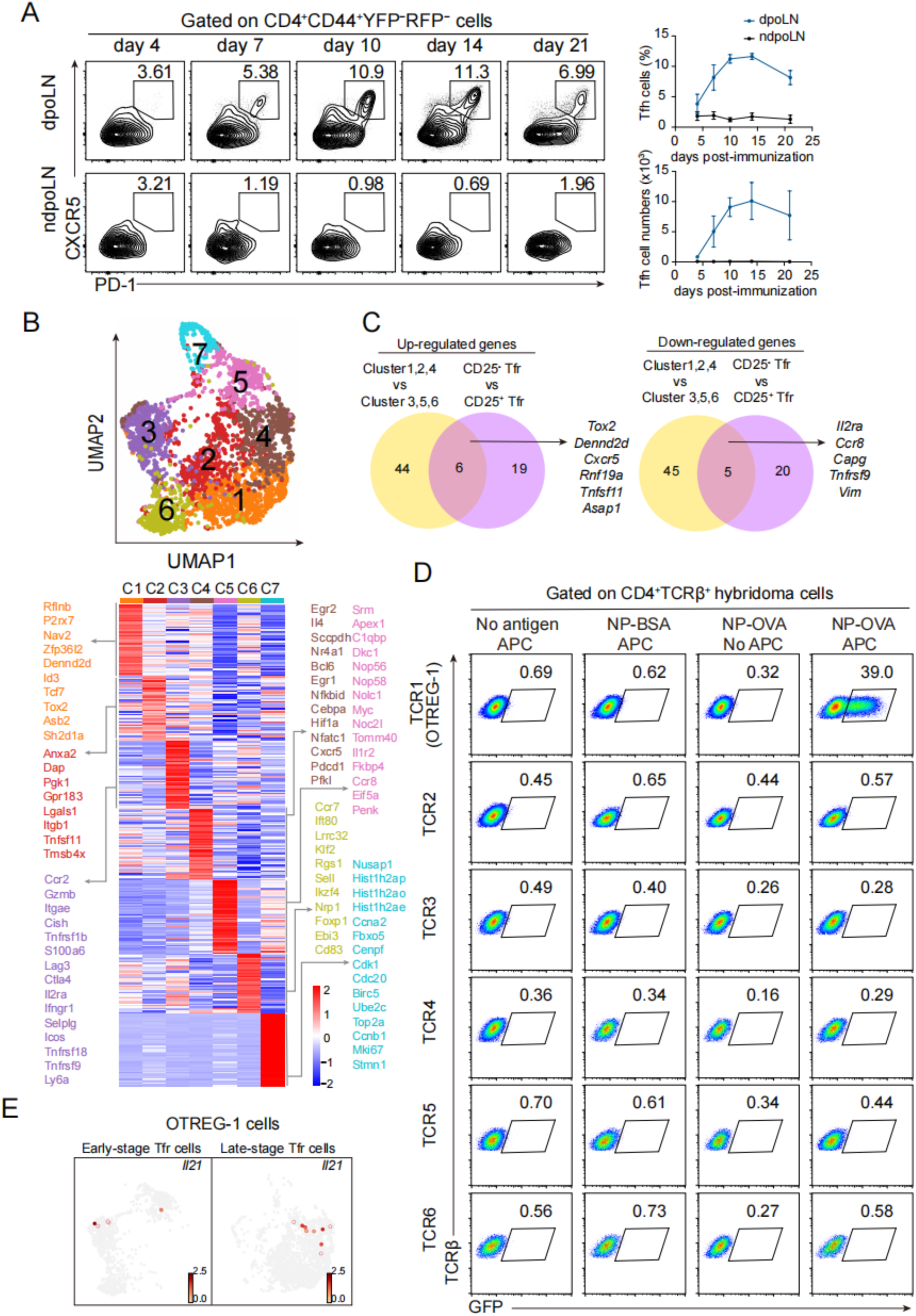
Single-cell transcriptomic profiling of T_FR_ cells. (**A**) Kinetics of T_FH_ cell differentiation post-immunization. (left) Flow cytometry plots showing PD-1 and CXCR5 expression on CD4^+^CD44^+^YFP^-^RFP^-^ T_FH_ cells in draining and non-draining popliteal lymph nodes (dpoLNs and ndpoLNs) at indicated time points. (right) Quantification of T_FH_ cell frequency (top) and absolute cell numbers (bottom). Day4 (n=3), day7 (n=5), day10 (n=4), day14 (n=3), day21(n=2). Flow cytometry data were acquired from two independent experiments. (**B**) Transcriptional signatures of T_FR_ clusters. Heatmap of differentially expressed genes (DEGs) across Tfr cell clusters (C1–C7). (**C**) Consensus gene signatures of late-stage T_FR_ cells. Venn diagram comparing DEGs between: Late-stage (C1/C2/C4) versus early-stage (C3/C5/C6) T_FR_ cell clusters; CD25^−^ versus CD25^+^ T_FR_ cells (**D**) Antigen-specific TCR activation assay. GFP induction in hybridomas expressing six expanded TCR clonotypes (OTREG-1, TCR2–6) after 48 h co-culture with: NP-OVA plus APCs (positive conditions), NP-OVA without APCs and NP-BSA or medium (negative controls). Representative flow cytometry data of two independent experiment. (**E**) UMAP highlighting the dominant OTREG-1 TCR clone with *Il21* expression.

**fig. S2.**
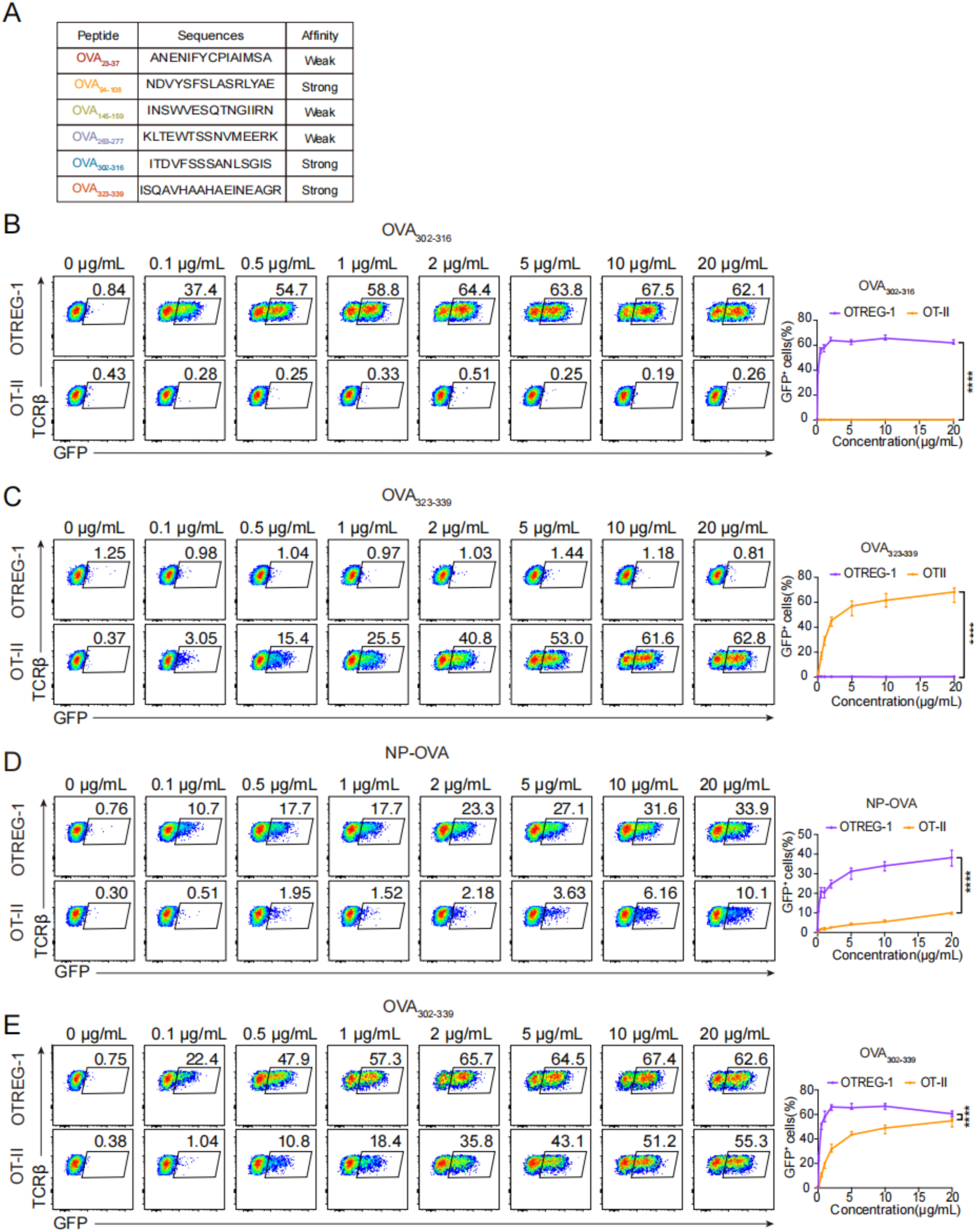
Distinct OVA epitope recognition by OTREG-1 and OT-II TCRs. (**A**) Predicted OVA peptide affinities. Structural modeling of six OVA-derived peptides ranked by estimated TCR-binding affinity. (**B**) Dose-response to OVA_302-316_ peptide. Percentage of GFP^+^ OTREG-1 (purple) and OT-II (orange) hybridoma cells after 48h coculture with indicated concentrations of OVA_302-316_ peptide. Data represent mean ± SD of triplicate experiments. (**C**) Dose-response to OVA_323-339_ peptide. Percentage of GFP^+^ cells for each hybridoma line cultured with graded doses of OVA_323-339_ peptide. Data represent mean ± SD of triplicate experiments. (**D**) Dose-response to NP-OVA. Percentage of GFP^+^ OTREG-1 (purple) and OT-II (orange) hybridoma cells after 48 h coculture with indicated concentrations of NP-OVA. Data represent mean ± SD of triplicate experiments. (**E**) Dose-response to OVA_302-339_. Percentage of GFP^+^ OTREG-1 (purple) and OT-II (orange) hybridoma cells after 48 h coculture with indicated concentrations of OVA_302-339_ peptide. Representative flow cytometry data of four independent experiment(left). Error bars show mean ± SD, *P* values were calculated using multiple unpaired Student’s *t*-test, \*\*\*\**P*<0.0001.

**fig. S3.**
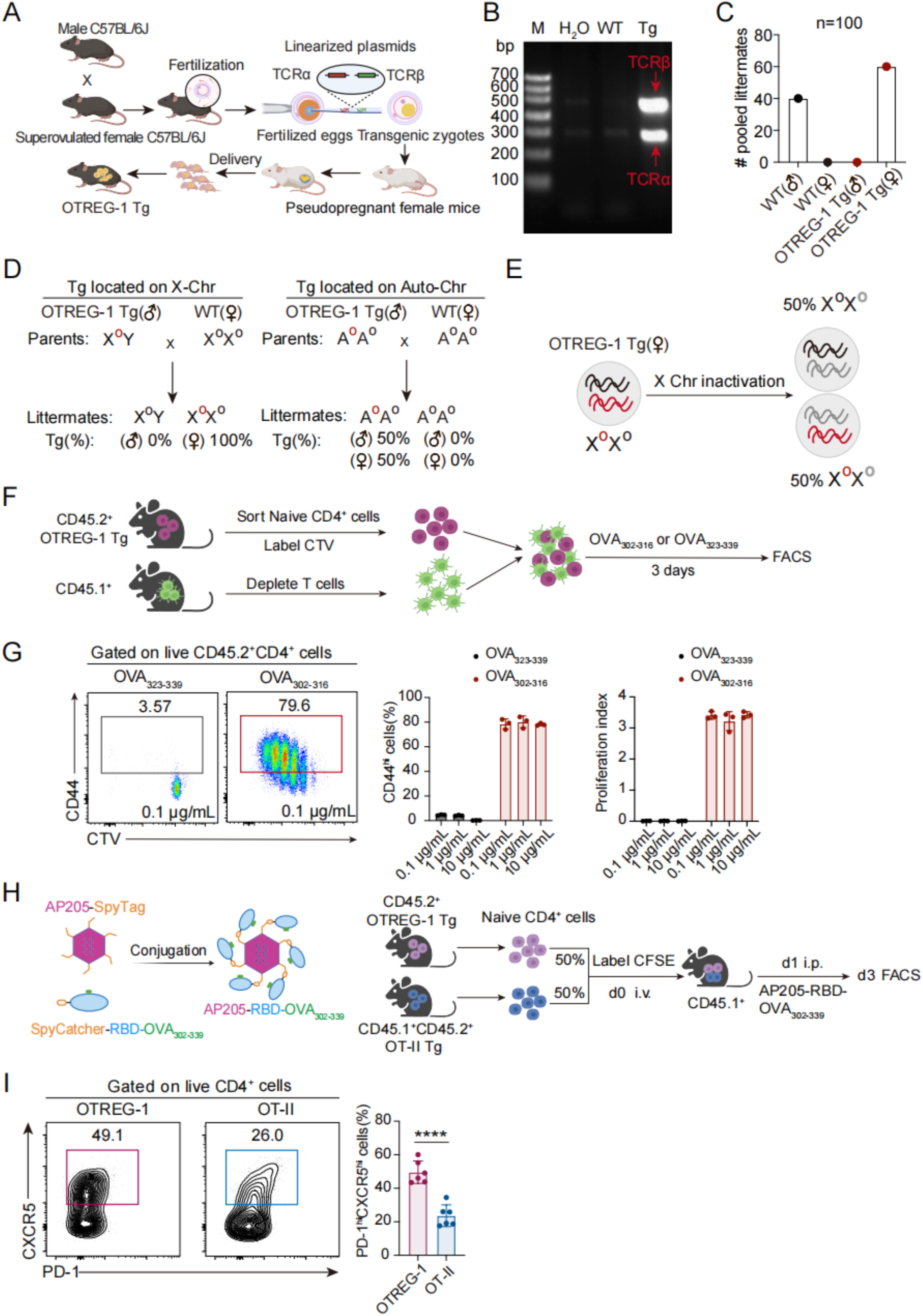
Validation of OVA-specific OTREG-1 transgenic mice. (**A**) OTREG-1 TCR Tg mouse generation. Schematic of the transgenic construct and breeding strategy to generate OTREG-1 TCR transgenic mice. (**B**) Genotype verification. Representative PCR showing TCRβ (517 bp) and TCRα (313 bp) transgene bands in WT versus OTREG-1 Tg mice. (**C**) Inheritance pattern analysis. Genotype and sex distribution of 100 offspring from male OTREG-1 Tg × female WT crosses, showing X-linked inheritance (all males WT, all females Tg). (**D**) Chromosomal localization models. Two potential transgene localization patterns: X-chromosome linkage (consistent with observed data) versus autosomal inheritance (inconsistent with observations). (**E**) X-chromosome inactivation. Theoretical expression pattern in female OTREG-1 Tg mice showing random X-inactivation (50% of cells express transgenic TCR). (**F**) *In vitro* validation assay. Experimental design: Naive CD4^+^CD25^-^CD44^lo^CD62L^hi^ T cells from male OTREG-1 Tg mice (CD45.2^+^) were CTV-labeled and co-cultured with WT APCs (CD45.1^+^) plus titrated OVA_302-316_ or OVA_323-339_ peptides (0.1-10 μg/mL) for 72h. (**G**) Antigen-specific activation *in vitro*. (left) Flow cytometry plots of CD44 expression and proliferation (CTV dilution) in CD45.2^+^CD4^+^ cells. (right) Quantification of CD44^hi^ cells and proliferation index. (**H**) *In vivo* particle antigen challenge. Construction of AP205-RBD-OVA_302-339_ nanoparticles (left). Competitive transfer experiment (right): CFSE-labeled 1:1 mix of naive OTREG-1 Tg (CD45.2^+^) and OT-II Tg (CD45.1^+^CD45.2^+^) cells transferred to WT hosts, followed by AP205-RBD-OVA_302-339_ immunization. (**I**) T_FH_ differentiation *in vivo*. (left) PD-1 and CXCR5 expression on donor cells. (right) Quantification of PD-1^hi^CXCR5^hi^ T_FH_ cells among OTREG-1 Tg (n=6) versus OT-II Tg (n=6) populations. All flow cytometry data were acquired from at least two independent experiments. Error bars show mean ± SD, *P* values were calculated using unpaired Student’s *t*-test, \*\*\*\**P*<0.0001.

**fig. S4.**
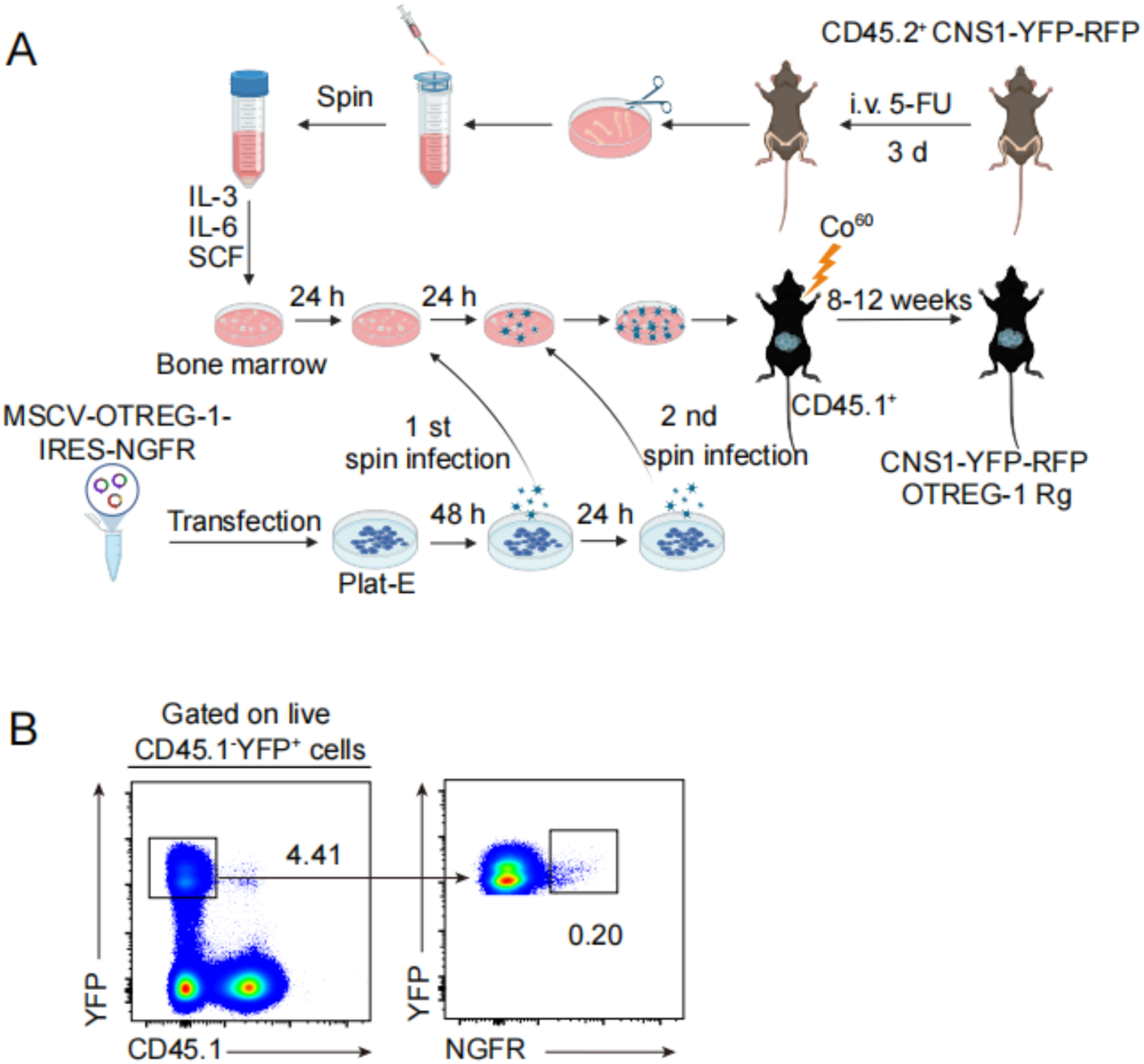
Generation of OVA-specific OTREG-1 retrogenic mice. (**A**) Schematic of OTREG-1 retrogenic mice generation. Bone marrow cells were isolated from 6–8 week-old CD45.2^+^ CNS1-YFP-RFP mice and infected with a retrovirus encoding the OTREG-1 TCRα and TCRβ chains. After transduction, the cells were intravenously transferred into irradiated CD45.1^+^ wildtype mice and allowed to reconstitute for 8-12 weeks. (**B**) The percentage of OTREG-1-expressing tT_reg_ cells (NGFR^+^) was shown in the OTREG-1 retrogenic mice after reconstitution.

**fig. S5.**
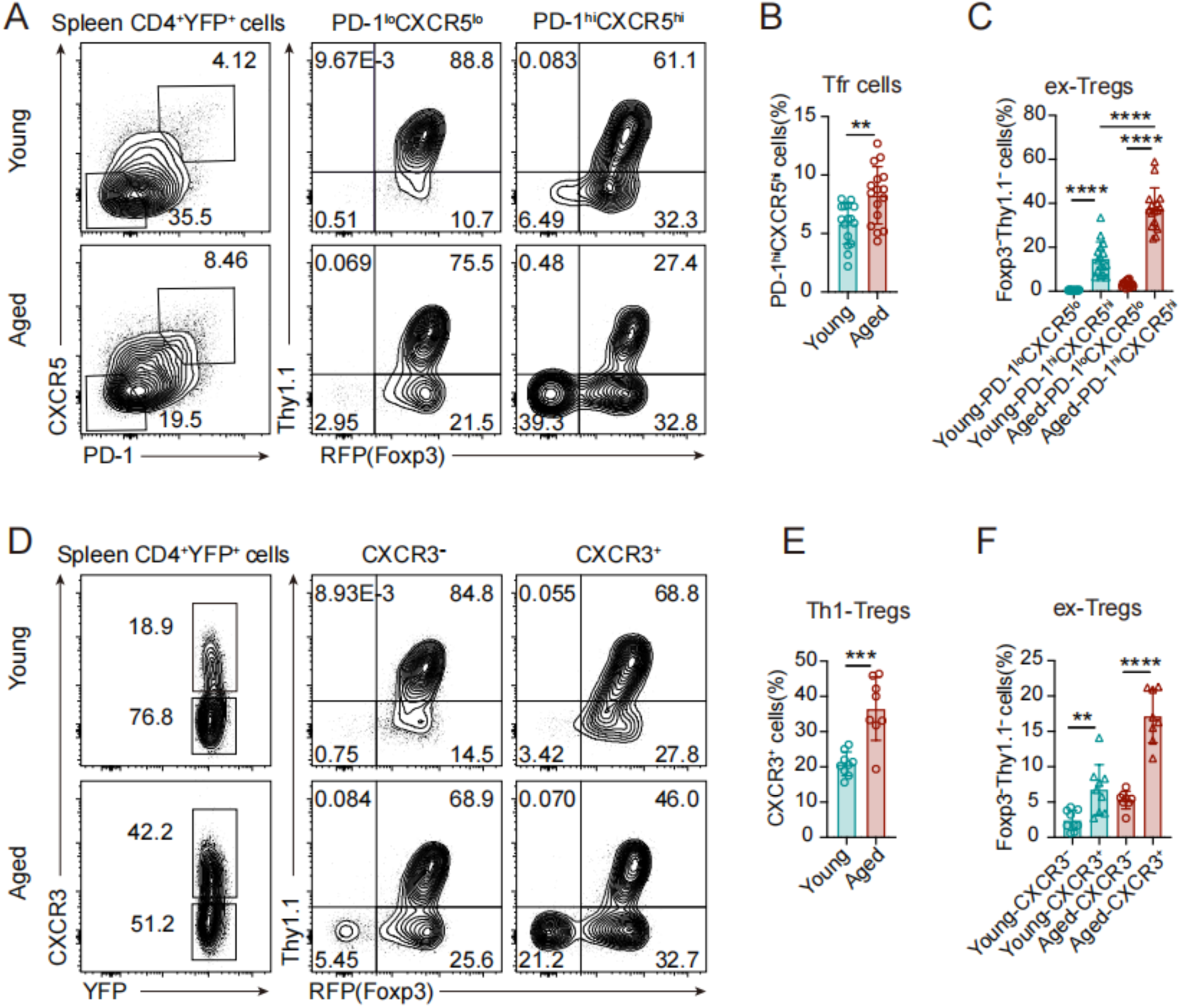
Age-associated increases in T_FR_/T_H_1-T_reg_ differentiation and Foxp3 instability. (**A**) Age-dependent T_FR_ cell accumulation and instability. Flow cytometry analysis of PD-1 and CXCR5 expression on CD4^+^YFP^+^ splenocytes from young (6–8 week) (n=9) versus aged (8–12 month) (n=8) CNS1-YFP-RFP mice (left). Foxp3 and Thy1.1 expression in PD-1^lo^CXCR5^lo^ versus PD-1^hi^CXCR5^hi^ subsets (right). (**B**) Quantification of T_FR_ cell expansion with age. Percentage of PD-1^hi^CXCR5^hi^ cells among CD4^+^YFP^+^ splenocytes. (**C**) Foxp3 instability in aged T_FR_ cells. Frequency of Foxp3^−^Thy1.1^−^ ex-T_reg_ cells in the PD-1^hi^CXCR5^hi^ subset. (**D**) T_H_1-T_reg_ cells differentiation and instability with aging. CXCR3 expression on CD4^+^YFP^+^ splenocytes (left). Foxp3 and Thy1.1 expression in CXCR3^−^ versus CXCR3^+^ subsets (right). (**E**) Age-related T_H_1-T_reg_ cells accumulation. Percentage of CXCR3^+^ cells among CD4^+^YFP^+^ splenocytes. (**F**) Foxp3 loss in T_H_1-T_reg_ subsets. Frequency of Foxp3^−^Thy1.1^−^ ex-T_reg_ cells in CXCR3^−^ versus CXCR3^+^ populations. All flow cytometry data were acquired from at least two independent experiments. Error bars show mean ± SD, *P* values were calculated using unpaired Student’s *t*-test, \*\**P*<0.01, \*\*\**P*<0.001, \*\*\*\**P*<0.0001.

## References

1. S. Sakaguchi, N. Mikami, J. B. Wing, A. Tanaka, K. Ichiyama, N. Ohkura, Regulatory T Cells and Human Disease. Annu. Rev. Immunol. 38, 541–566 (2020).

2. M. E. Brunkow, E. W. Jeffery, K. A. Hjerrild, B. Paeper, L. B. Clark, S. A. Yasayko, J. E. Wilkinson, D. Galas, S. F. Ziegler, F. Ramsdell, Disruption of a new forkhead/winged-helix protein, scurfin, results in the fatal lymphoproliferative disorder of the scurfy mouse. Nat. Genet. 27, 68–73 (2001).

3. R. S. Wildin, F. Ramsdell, J. Peake, F. Faravelli, J. L. Casanova, N. Buist, E. Levy-Lahad, M. Mazzella, O. Goulet, L. Perroni, F. D. Bricarelli, G. Byrne, M. McEuen, S. Proll, M. Appleby, M. E. Brunkow, X-linked neonatal diabetes mellitus, enteropathy and endocrinopathy syndrome is the human equivalent of mouse scurfy. Nat. Genet. 27, 18–20 (2001).

4. L. M. Williams, A. Y. Rudensky, Maintenance of the Foxp3-dependent developmental program in mature regulatory T cells requires continued expression of Foxp3. Nat Immunol. 8, 277–284 (2007).

5. Y. Y. Wan, R. A. Flavell, Regulatory T-cell functions are subverted and converted owing to attenuated Foxp3 expression. Nature. 445, 766–770 (2007).

6. A. G. Levine, A. Arvey, W. Jin, A. Y. Rudensky, Continuous requirement for the TCR in regulatory T cell function. Nat Immunol. 15, 1070–1078 (2014).

7. Y. Chung, S. Tanaka, F. Chu, R. I. Nurieva, G. J. Martinez, S. Rawal, Y. H. Wang, H. Lim, J. M. Reynolds, X. H. Zhou, H. M. Fan, Z. M. Liu, S. S. Neelapu, C. Dong, Follicular regulatory T cells expressing Foxp3 and Bcl-6 suppress germinal center reactions. Nat Med. 17, 983–988 (2011).

8. M. A. Linterman, W. Pierson, S. K. Lee, A. Kallies, S. Kawamoto, T. F. Rayner, M. Srivastava, D. P. Divekar, L. Beaton, J. J. Hogan, S. Fagarasan, A. Liston, K. G. Smith, C. G. Vinuesa, Foxp3+ follicular regulatory T cells control the germinal center response. Nat Med. 17, 975–982 (2011).

9. M. A. Koch, G. Tucker-Heard, N. R. Perdue, J. R. Killebrew, K. B. Urdahl, D. J. Campbell, The transcription factor T-bet controls regulatory T cell homeostasis and function during type 1 inflammation. Nat Immunol. 10, 595–602 (2009).

10. M. DuPage, J. A. Bluestone, Harnessing the plasticity of CD4(+) T cells to treat immune-mediated disease. Nat Rev Immunol. 16, 149–163 (2016).

11. Z. Zhang, W. Zhang, J. Guo, Q. Gu, X. Zhu, X. Zhou, Activation and Functional Specialization of Regulatory T Cells Lead to the Generation of Foxp3 Instability. J Immunol. 198, 2612–2625 (2017).

12. X. Zhou, S. L. Bailey-Bucktrout, L. T. Jeker, C. Penaranda, M. Martinez-Llordella, M. Ashby, M. Nakayama, W. Rosenthal, J. A. Bluestone, Instability of the transcription factor Foxp3 leads to the generation of pathogenic memory T cells in vivo. Nat Immunol. 10, 1000–1007 (2009).

13. M. E. Birnbaum, J. L. Mendoza, D. K. Sethi, S. Dong, J. Glanville, J. Dobbins, E. Ozkan, M. M. Davis, K. W. Wucherpfennig, K. C. Garcia, Deconstructing the peptide-MHC specificity of T cell recognition. Cell. 157, 1073–1087 (2014).

14. A. K. Sewell, Why must T cells be cross-reactive? Nat Rev Immunol. 12, 669–677 (2012).

15. A. R. Maceiras, S. C. P. Almeida, E. Mariotti-Ferrandiz, W. Chaara, F. Jebbawi, A. Six, S. Hori, D. Klatzmann, J. Faro, L. Graca, T follicular helper and T follicular regulatory cells have different TCR specificity. Nat Commun. 8, 15067 (2017).

16. M. Palatella, S. M. Guillaume, M. A. Linterman, J. Huehn, The dark side of Tregs during aging. Front Immunol. 13, 940705 (2022).

17. S. Ma, Z. Ji, B. Zhang, L. Geng, Y. Cai, C. Nie, J. Li, Y. Zuo, Y. Sun, G. Xu, B. Liu, J. Ai, F. Liu, L. Zhao, J. Zhang, H. Zhang, S. Sun, H. Huang, Y. Zhang, Y. Ye, Y. Fan, F. Zheng, J. Hu, B. Zhang, J. Li, X. Feng, F. Zhang, Y. Zhuang, T. Li, Y. Yu, Z. Bao, S. Pan, C. Rodriguez Esteban, Z. Liu, H. Deng, F. Wen, M. Song, S. Wang, G. Zhu, J. Yang, T. Jiang, W. Song, J. C. Izpisua Belmonte, J. Qu, W. Zhang, Y. Gu, G. H. Liu, Spatial transcriptomic landscape unveils immunoglobin-associated senescence as a hallmark of aging. Cell. 187, 7025–7044 e7034 (2024).

18. X. Liu, Z. Liu, Z. Wu, J. Ren, Y. Fan, L. Sun, G. Cao, Y. Niu, B. Zhang, Q. Ji, X. Jiang, C. Wang, Q. Wang, Z. Ji, L. Li, C. R. Esteban, K. Yan, W. Li, Y. Cai, S. Wang, A. Zheng, Y. E. Zhang, S. Tan, Y. Cai, M. Song, F. Lu, F. Tang, W. Ji, Q. Zhou, J. C. I. Belmonte, W. Zhang, J. Qu, G. H. Liu, Resurrection of endogenous retroviruses during aging reinforces senescence. Cell. 186, 287–304 e226 (2023).

19. J. Merkenschlager, S. Finkin, V. Ramos, J. Kraft, M. Cipolla, C. R. Nowosad, H. Hartweger, W. Zhang, P. D. B. Olinares, A. Gazumyan, T. Y. Oliveira, B. T. Chait, M. C. Nussenzweig, Dynamic regulation of T(FH) selection during the germinal centre reaction. Nature. 591, 458–463 (2021).

20. Y. Lin, Z. Wan, B. Liu, J. Yao, T. Li, F. Yang, J. Sui, Y. Zhao, W. Liu, X. Zhou, J. Wang, H. Qi, B cell-reactive triad of B cells, follicular helper and regulatory T cells at homeostasis. Cell Res. 34, 295–308 (2024).

21. J. B. Wing, Y. Kitagawa, M. Locci, H. Hume, C. Tay, T. Morita, Y. Kidani, K. Matsuda, T. Inoue, T. Kurosaki, S. Crotty, C. Coban, N. Ohkura, S. Sakaguchi, A distinct subpopulation of CD25(-) T-follicular regulatory cells localizes in the germinal centers. Proc Natl Acad Sci U S A. 114, E6400–E6409 (2017).

22. H. Kishimoto, J. Sprent, Negative selection in the thymus includes semimature T cells. J Exp Med. 185, 263–271 (1997).

23. C. Guo, Y. Peng, L. Lin, X. Pan, M. Fang, Y. Zhao, K. Bao, R. Li, J. Han, J. Chen, T. Z. Song, X. L. Feng, Y. Zhou, G. Zhao, L. Zhang, Y. Zheng, P. Zhu, H. Hang, L. Zhang, Z. Hua, H. Deng, B. Hou, A pathogen-like antigen-based vaccine confers immune protection against SARS-CoV-2 in non-human primates. Cell Rep Med. 2, 100448 (2021).

24. W. B. Zhang, F. Sun, D. A. Tirrell, F. H. Arnold, Controlling macromolecular topology with genetically encoded SpyTag-SpyCatcher chemistry. J. Am. Chem. Soc. 135, 13988–13997 (2013).

25. K. Luthje, A. Kallies, Y. Shimohakamada, G. T. Belz, A. Light, D. M. Tarlinton, S. L. Nutt, The development and fate of follicular helper T cells defined by an IL-21 reporter mouse. Nat Immunol. 13, 491–498 (2012).

26. R. Wang, Q. Wan, L. Kozhaya, H. Fujii, D. Unutmaz, Identification of a regulatory T cell specific cell surface molecule that mediates suppressive signals and induces Foxp3 expression. PLoS ONE. 3, e2705 (2008).

27. K. Schumann, S. S. Raju, M. Lauber, S. Kolb, E. Shifrut, J. T. Cortez, N. Skartsis, V. Q. Nguyen, J. M. Woo, T. L. Roth, R. Yu, M. L. T. Nguyen, D. R. Simeonov, D. N. Nguyen, S. Targ, R. E. Gate, Q. Tang, J. A. Bluestone, M. H. Spitzer, C. J. Ye, A. Marson, Functional CRISPR dissection of gene networks controlling human regulatory T cell identity. Nat Immunol. 21, 1456–1466 (2020).

28. R. I. Nurieva, Y. Chung, G. J. Martinez, X. O. Yang, S. Tanaka, T. D. Matskevitch, Y. H. Wang, C. Dong, Bcl6 mediates the development of T follicular helper cells. Science. 325, 1001–1005 (2009).

29. X. Wu, Y. Wang, R. Huang, Q. Gai, H. Liu, M. Shi, X. Zhang, Y. Zuo, L. Chen, Q. Zhao, Y. Shi, F. Wang, X. Yan, H. Lu, S. Xu, X. Yao, L. Chen, X. Zhang, Q. Tian, Z. Yang, B. Zhong, C. Dong, Y. Wang, X. W. Bian, X. Liu, SOSTDC1-producing follicular helper T cells promote regulatory follicular T cell differentiation. Science. 369, 984–988 (2020).

30. W. Xu, X. Zhao, X. Wang, H. Feng, M. Gou, W. Jin, X. Wang, X. Liu, C. Dong, The Transcription Factor Tox2 Drives T Follicular Helper Cell Development via Regulating Chromatin Accessibility. Immunity. 51, 826–839 e825 (2019).

31. X. Tai, A. Indart, M. Rojano, J. Guo, N. Apenes, T. Kadakia, M. Craveiro, A. Alag, R. Etzensperger, M. E. Badr, F. Zhang, Z. Zhang, J. Mu, T. Guinter, A. Crossman, L. Granger, S. Sharrow, X. Zhou, A. Singer, How autoreactive thymocytes differentiate into regulatory versus effector CD4(+) T cells after avoiding clonal deletion. Nat Immunol. 24, 637–651 (2023).

32. L. Klein, E. A. Robey, C. S. Hsieh, Central CD4(+) T cell tolerance: deletion versus regulatory T cell differentiation. Nat Rev Immunol. 19, 7–18 (2019).

33. K. S. Smigiel, E. Richards, S. Srivastava, K. R. Thomas, J. C. Dudda, K. D. Klonowski, D. J. Campbell, CCR7 provides localized access to IL-2 and defines homeostatically distinct regulatory T cell subsets. J Exp Med. 211, 121–136 (2014).

34. W. Ouyang, W. Liao, C. T. Luo, N. Yin, M. Huse, M. V. Kim, M. Peng, P. Chan, Q. Ma, Y. Mo, D. Meijer, K. Zhao, A. Y. Rudensky, G. Atwal, M. Q. Zhang, M. O. Li, Novel Foxo1-dependent transcriptional programs control T(reg) cell function. Nature. 491, 554–559 (2012).

35. Z. Zhang, X. Zhou, Foxp3 Instability Helps tTregs Distinguish Self and Non-self. Front Immunol. 10, 2226 (2019).

36. S. Srinivas, T. Watanabe, C. S. Lin, C. M. William, Y. Tanabe, T. M. Jessell, F. Costantini, Cre reporter strains produced by targeted insertion of EYFP and ECFP into the ROSA26 locus. BMC Dev. Biol. 1, 4 (2001).

37. M. Xu, M. Pokrovskii, Y. Ding, R. Yi, C. Au, O. J. Harrison, C. Galan, Y. Belkaid, R. Bonneau, D. R. Littman, c-MAF-dependent regulatory T cells mediate immunological tolerance to a gut pathobiont. Nature. 554, 373–377 (2018).

38. T. Wang, J. Guo, L. Liping, Q. Jin, F. Zhang, B. Hou, Y. Zhang, X. Zhou, The histone lysine methyltransferase MLL1 regulates the activation and functional specialization of regulatory T cells. Cell Rep. 43, 114222 (2024).

39. D. W. Cain, S. E. Sanders, M. M. Cunningham, G. Kelsoe, Disparate adjuvant properties among three formulations of "alum". Vaccine. 31, 653–660 (2013).

